# Adaptive benefits of motility in cross-feeding mutualisms

**DOI:** 10.64898/2026.08.14.744873

**Authors:** Naven Narayanan Venkatanarayanan, Jonathan N.V. Martinson, Allison K. Shaw, William R. Harcombe

## Abstract

Cross-feeding mutualisms, in which partner species exchange essential metabolites, are ubiquitous in microbial communities. In spatially structured environments, motility can improve access to partner-produced resources but also impose metabolic costs and displace cells from nutrient-rich regions, so its net benefit depends on the spatial dynamics of the interaction. Here, we combine competition experiments in a cross-feeding mutualism between *Escherichia coli* and *Salmonella enterica* with a spatially explicit consumer-resource model to determine what drives selection on motility. Spatial structure imposes asymmetric selection between partners i.e. *S. enterica* benefits from motility regardless of partner motility, whereas selection on *E. coli* switches from favourable to unfavourable depending on whether its partner can move. Competition in well-mixed culture suggests that this reversal reflects the loss of a spatial benefit rather than an increased cost. Our model attributes the asymmetry to three interacting factors: the ratio of metabolite production to consumption which sets whether the cross-fed resource is scarce or abundant; the number of growth-limiting resources which determines whether an alternative gradient can rescue the benefit of motility; and partner motility and growth rate, which shape where metabolites are produced. When a metabolite is scarce, motile cells gain by dispersing into regions it has reached but not yet been depleted from. When it accumulates, this gradient is eroded, and the motility costs offset any benefit it provides. Selection on motility therefore depends on the metabolic structure of the interaction and the spatial behaviour of partners.

## Introduction

Spatial structure is a fundamental property of ecological systems, shaping the organization and dynamics of complex communities across scales. In microbial communities especially, an individual’s location determines the abiotic environment it experiences (for e.g. pH, nutrient concentrations) and the neighbours it interacts with. Although often studied in well-mixed cultures, microbes in nature are predominantly found attached to surfaces or embedded in biofilms [1–3]. These local interactions with abiotic and biotic factors collectively influence individual growth dynamics and give rise to emergent eco-evolutionary patterns at the population and community scales [4–6]. Understanding how spatial heterogeneity governs these dynamics is therefore crucial to predicting the coexistence, assembly, and function of microbial communities [7].

Interactions between individuals in microbial communities are often environmentally mediated such as competition for limiting resources, toxin production that suppresses competitor growth, or excreted metabolites that fuel the growth of neighbouring species [8–11]. The last of these, termed cross-feeding, is ubiquitous across the tree of life, occurring not only between bacterial species but also between bacteria and fungi, animals, protists, and plants [12–15]. Species engaged in cross-feeding reshape the local resource landscape through metabolite excretion, generating spatial heterogeneity that can drive selection on ecologically important traits.

One such trait is motility. In spatio-temporally varying environments, motility allows individuals to exploit resource-rich regions, conferring an apparent growth advantage [16–18]. Yet motility also carries substantial costs such as flagellar construction and maintenance demand energy and the length scales over which metabolites diffuse may be short enough that movement offers limited benefit [19–23]. Indeed, recent work analysing over 11,000 bacterial genomes found that motility was lost far more than gained throughout evolutionary history suggesting that it can be beneficial but is not essential for survival [24]. Theoretical models of dispersal evolution predict that movement should generally be disfavoured in obligate mutualisms, because individuals that disperse away from their partners lose access to essential benefits [25–27]. However, these models typically assume spatially localised benefits, appropriate for contact-dependent interactions but potentially misleading for cross-feeding, where benefits are mediated by diffusible metabolites. Recent experimental work supports this distinction wherein random motility enhances metabolic coupling in spatially structured cross-feeding consortia by disrupting clonal clusters and increasing partner proximity [28]. Motility also provides a fitness advantage in cross-feeding communities in turbulent well-mixed environments [29]. Further, experimental evolution under cross-feeding preserves motility that might otherwise be lost when the same resource is supplied externally. When metabolites can spread through the environment, the cost of moving away from a partner may be offset by the ability to track where resources are being produced. Whether this distinction matters and how it interacts with the metabolic details of the interaction remains understudied. More broadly, how the spatial dynamics of metabolite exchange shape motility selection in mutualistic communities remains an open question, despite growing evidence that spatial structure fundamentally alters the ecological dynamics of cross-feeding interactions [30, 31].

Here, we address this question using a combination of theory and experiments. We leverage a well-established, stable, obligate cross-feeding mutualism between *E. coli* and *S. enterica* to study the conditions under which motile strains of a focal species are selected for or against in the presence of their partner. We then construct a resource-explicit model of two mutualistically interacting species in a spatially structured environment, identify the conditions under which motility is beneficial, and use the model to propose mechanisms driving selection for or against motility. We find that selection on motility can be predicted from three features of the interaction. First, the ratio of metabolite production to consumption, which determines whether the cross-fed resource is scarce or abundant. Second, the number of resources limiting a species’ growth. And third, the partner’s motility status, which shapes the spatial distribution of metabolite production. These results reveal how spatial feedbacks between mutualism and movement shape the evolution of motility across spatially structured habitats.

## Methods

### Biological system

We study a well-characterised cross-feeding mutualism between *Escherichia coli* and *Salmonella enterica* [32, 33]. When grown on lactose as the sole carbon source, *E. coli* consumes lactose and excretes acetate and other carbon byproducts. *S. enterica* cannot utilize lactose but can grow on the excreted acetate. In return, *S. enterica* produces methionine, an essential amino acid that the *E. coli* strain used here cannot synthesize (Figure 1a). This reciprocal exchange of metabolites creates an obligate mutualism: neither species can grow in isolation under these conditions. Both species can exist as motile or non-motile strains, allowing us to ask how the spatial dynamics of cross-feeding shape selection on motility.

**Figure 1:**
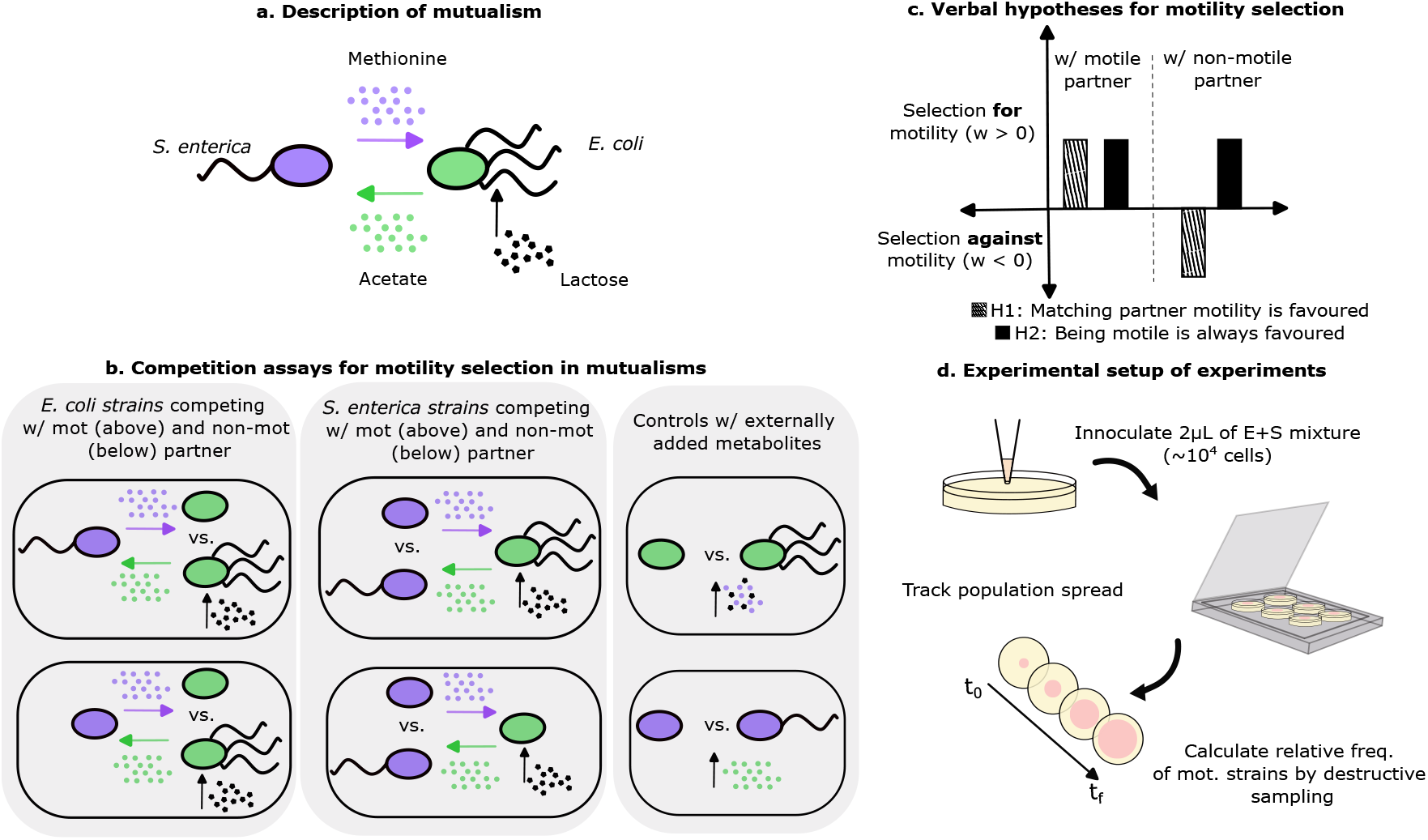
System description, hypotheses, and experimental setup: (a) shows a schematic of the experimental cross-feeding mutualisms with *E. coli* and *S. enterica* with the relevant metabolite exchanges. (b) shows the experimental treatments of competition between motile and non-motile strains of a focal species in the presence of their mutualistic partner. (c) shows competing biological hypotheses predicting selection outcomes for motility with partners of differing motility status. (d) shows how spreading of bacterial colonies in which changes in mutant (non-motile) frequency was tracked over time following inoculation of community at the centre of the petridish.

### Experimental Methods

The *E. coli* strain used in this study is an MG1655 derivative containing a Δ*metB* mutation. This strain was established as the motile ‘E’ strain. To this strain, we performed a transduction with P1 phage to generate a Δ*fliC* mutant with antibiotic resistance to kanamycin [34, 35]. This strain was defined as the ‘E’ non-motile strain. To build the *Salmonella* knockout we used P22 HT int transduction to move the *flgE* knockout from the BEI Resources *S. enterica* 14028s knockout library into our strain [36]. The antibiotic resistance in non-motile strains was used to determine relative frequency of strains at the beginning versus the end of the experiment to estimate selection on motility.

The strains were cultured overnight in Lysogeny Broth (LB) with appropriate antibiotic selection at 37ºC with shaking. The next day, they were diluted into the same broth, grown at 37ºC until they reached mid-log phase, and washed and adjusted to OD600 = 0.2 in sterile saline (0.9% w/v). Washed and adjusted cells were serially diluted and plated onto LB agar to determine the initial population size of each strain. Co-cultures were grown in lactose Hypho minimal medium (2.92 mM lactose, 14.53 mM K_2_HPO_4_, 18.75 mM NaH_2_PO_4_, 3.78 mM (NH_4_)_2_SO_4_, 0.81 mM MgSO_4_, 20 *µ*M CaCl_2_, and trace metals) solidified with 0.3% (w/v) agar for swimming assays or 1.5% (w/v) agar for plating. Monocultures of *E. coli* were supplemented with 50 *µ*M L-methionine. Monocultures of *S. enterica* replaced lactose with 5.84 mM galactose as the sole carbon source.

#### Swimming experiments

In these experiments, strains from the focal species (either *E. coli* or *S. enterica*) were competed at a 90:10 starting ratio of motile to non-motile individuals. In reality, the experimental initial ratios deviated slightly from the 90:10 ratio (see Figure S2). In the mutualistic co-culture experiments, the partner species’ population size was the same as the focal species. The focal species strains and, where appropriate, the partner species were mixed. Next, either 1µL or 2µL of cell mixtures was inoculated into the centre of a swimming agar plate for the monoculture and co-culture experiments, respectively. For each species, roughly 10^4^ cells were inoculated into swimming agar (0.3 % w/v). The inoculated plates were incubated at 37º C in a well-humidified incubator for five days (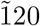 hours). If the swimming cells reached the edge of the plate, the experiment was ended for that treatment to avoid overgrowth. The different experimental (and in-silico) treatments are described in a schematic in Figure 1b. In our experiments, we focused on the swimming behaviour of the bacterial strains as opposed to surface level swarming or growth mediated expansion (found on hard agar surfaces). Thus our inoculation was performed by gently pricking the surface of the swimming agar to introduce the cells below the agar surface.

Following spatial expansion, to harvest cells, a 50 mL serological pipette was used to aspirate all of the agar in the petridish, then dispensed into a 50 mL centrifuge tube. The weight of the aspirated agar was measured by weighing the tubes before and after the addition of agar. The aspirated agar was vortexed for ≈ 30 seconds until it became a slurry. The slurry was serially diluted and plated onto selective solid agar with a multichannel pipette. Quantification plates were incubated for 24-48 hours at 37ºC and colonies were counted.

#### Liquid culture experiments to determine cost of motility

In order to ascertain whether there were any costs associated with having a motile apparatus (for instance, flagella), we performed the same competition assay as described above in liquid media. The assay was performed in 200 *µ*L of well-mixed liquid medium for 24 h. In this assay, the mutualism benefits exchanged (i.e. acetate and methionine) are accessible equally by both the motile and non-motile strains of the focal species due to the lack of spatial structure. By removing spatial structure, we largely eliminate the benefit of motility and study the effects of having a motile structure on strain growth rate.

### Data Analysis

Data analysis was performed in R 4.5.3 using custom scripts available at github.com/naven22/motility_evolution_mutualism. Briefly, fitness was calculated as *w* = − log_2_ (*f*_non,*f*_ */f*_non,0_), where *f*_non_ is the frequency of the non-motile strain of the focal species. All selection coefficients in this paper, from both experiments and simulations, are reported in log_2_ units, so that *w* = 1 corresponds to a doubling of the motile strain’s relative representation. Neutral fitness is centred on zero, values below zero indicate selection against motility, and values above zero indicate selection for motility. Selection coefficients were tested against zero using one-sample two-tailed *t*-tests (*n* = 5 biological replicates, df = 4) with Holm-Bonferroni correction applied across the six treatments in each assay. Differences between partner conditions were assessed with Tukey’s HSD.

Because *w* is cumulative, it scales with the number of generations over which selection acts, and the two assays differ both in duration (24 h in liquid, five days on agar) and in inoculum dilution (2 *µ*L into a 200 *µ*L well versus a 90mm agar plate). We therefore also express selection per generation. Total population size at the endpoint was calculated as the plated colony count converted to CFU mL^−1^ and multiplied by the culture volume which was 0.2 mL for liquid cultures, and for swimming plates the mass of aspirated agar which was at 0.3% (w/v). The number of focal-population doublings is

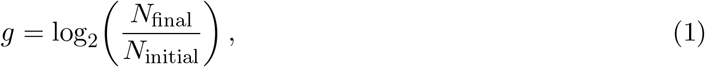

and per-generation selection is *s* = *w/g*. We calculate the generation counts for liquid and swimming experiments in the SI (see Table S4).

As detailed in the earlier sections, the liquid and agar assays measure different quantities. In liquid culture there is no spatial structure, thus exchanged metabolites are equally accessible to motile and non-motile cells and the primary fitness consequence of motility is the metabolic cost of the flagellar apparatus. On swimming agar, the measured selection reflects that cost together with any spatial benefit of motility itself. We can therefore decompose selection (per generation) as

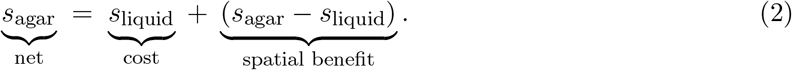

The two assays are independent experiments with unpaired replicates, so the spatial benefit is a difference of independent means with its significance assessed by Welch’s *t*-test with Holm-Bonferroni correction across the six treatments.

### Mathematical model

To understand how partner motility shapes selection on motility, we developed a spatially explicit model of the *E. coli* -*S. enterica* interaction. Both species exist as motile and non-motile strains, where motile cells spread faster but pay a fractional growth rate cost *c*. We modelled population abundance and resource concentration dynamics using reaction-diffusion equations in one spatial dimension. For *E. coli* :

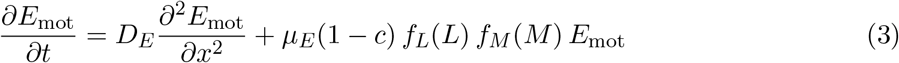

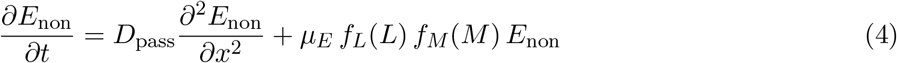

and for *S. enterica*:

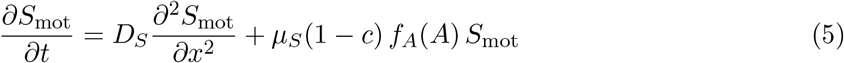

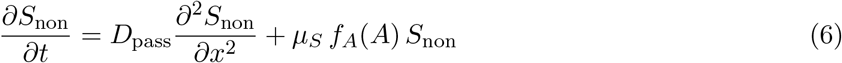

where *D*_*E*_ and *D*_*S*_ are diffusion coefficients for motile cells, *D*_pass_ is a much smaller passive diffusion coefficient for non-motile cells, and *µ*_*E*_ and *µ*_*S*_ are maximum growth rates. Growth limitation follows Monod kinetics, *f*(*R*) = *R/*(*K* + *R*), where *R* is resource concentration and *K* is the half-saturation constant. *S. enterica* growth depends on acetate (*A*) alone. ‘c’ varies from 0 i.e. no motility cost to 1, high cost resulting in zero growth rate.

Resource dynamics couple the two species through cross-feeding. Consumption and production are proportional to the realised growth rate of each strain, so that a motile cell, which divides at a rate reduced by the factor (1 − *c*), likewise consumes and produces at a correspondingly reduced rate. We therefore define the growth-weighted densities

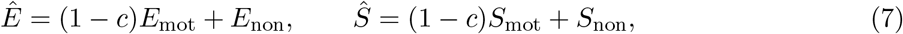

and the resource dynamics are

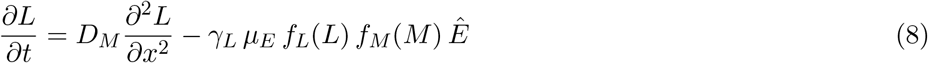

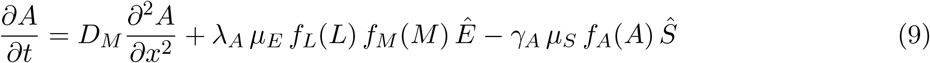

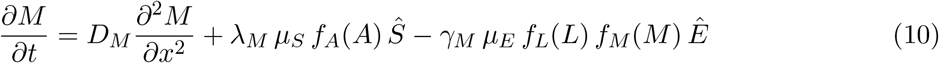

where *D*_*M*_ is the metabolite diffusion coefficient, *λ* denotes production yields (g metabolite per cell division) and *γ* denotes consumption yields. This formulation ensures that metabolite flux is strictly proportional to growth, so that non-growing cells neither consume nor produce resources and the motility cost *c* acts on growth rate alone rather than imposing a second, implicit penalty on yield. The ratios *ρ*_*M*_ = *λ*_*M*_ */γ*_*M*_ and *ρ*_*A*_ = *λ*_*A*_*/γ*_*A*_ describe the production-to-consumption balance of each cross-fed metabolite. Because each species’ production of one metabolite is fuelled by its consumption of the other, these ratios do not act independently. At steady state, a closed cycle of exchange is self-sustaining only when

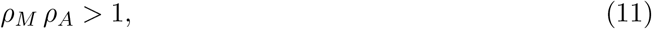

that is, when each unit of methionine consumed by *E. coli* ultimately returns more than one unit of methionine through the acetate it produces. When *ρ*_*M*_ *ρ*_*A*_ < 1 the metabolites are exhausted and the mutualism is not self-sustaining. We also assume no-flux boundary conditions to ensure a closed system across the entire range of simulations. We highlight that Eq. 11 constrains the product of the two accumulation ratios, which governs whether the exchange cycle is viable. Whether a given metabolite is locally scarce or abundant for a species is governed by the individual ratio *ρ*_*A*_ or *ρ*_*M*_ of the metabolite that species receives. Full details of the model, non-dimensionalisation, and numerical implementation are provided in the SI. Our model assumes a single spatial dimension, whereas the experiments involve radial expansion on 2D agar surfaces. Because we model undirected (diffusive) motility, we note that it is isotropic by assumption, and the qualitative dynamics of spatial strain competition will be preserved in 1D.

We non-dimensionalised the model to identify key parameter groupings (see SI). The critical dimensionless parameters include *β*_*E*_ = *D*_*E*_*/D*_*M*_ (for *E. coli* ; a similar *β*_*S*_ can be written for *S. enterica*), the ratio of cell motility to metabolite diffusion, and Da = *D*_*M*_ */*(*µ*_*E*_ℒ^2^), which compares the timescale of metabolite diffusion to cell growth across the domain length ℒ. When *β* is small, metabolites spread faster than cells, reducing the importance of cell position for nutrient access.

Our simulations mirrored the four experimental treatments. The focal species comprised competing motile and non-motile strains (initially 90% and 10% respectively), while the partner species was fixed as entirely motile or entirely non-motile. This yielded four treatments: *E. coli* focal with motile *S. enterica* partner (E co Smot), *E. coli* focal with non-motile partner (E co Snon), *S. enterica* focal with motile *E. coli* partner (S co Emot), and *S. enterica* focal with non-motile partner (S co Enon). Both populations were initialised as Gaussian distributions centred in the domain, with cross-fed metabolites localised to the initial population distribution.

Selection was quantified as for the experiments, using non-motile frequencies integrated across space, enabling direct comparison of model and experiment. All simulations were run on MATLAB v. 2025a (scripts available at github.com/naven22/motility_evolution_mutualism). We ran each of our simulations to steady state and focus our interpretation on the lactose-limited regime (unshaded regions in Figure 5), as lactose is the primary carbon source in our experimental system and its depletion defines the natural endpoint of growth. We note that if one of the cross-fed metabolites is the first to get exhausted, then it implies a breakdown in the mutualism between the two species (shaded in grey; Figure 5).

## Results

### Partner motility determines selection on motility in *E. coli*

Competing motile against non-motile *E. coli* revealed that selection on motility depended on the motility status of the *S. enterica* partner (Figure 2a). With a motile partner, motile *E. coli* increased in frequency (*w* = 0.84 ± 0.23, *p* = 0.043, *n* = 5). With a non-motile partner the pattern reversed and non-motile *E. coli* was favoured (*w* = −0.88 ± 0.16, *p* = 0.014), a significant difference between treatments (Tukey HSD: Δ*w* = 1.720, *p* < 0.001).

**Figure 2:**
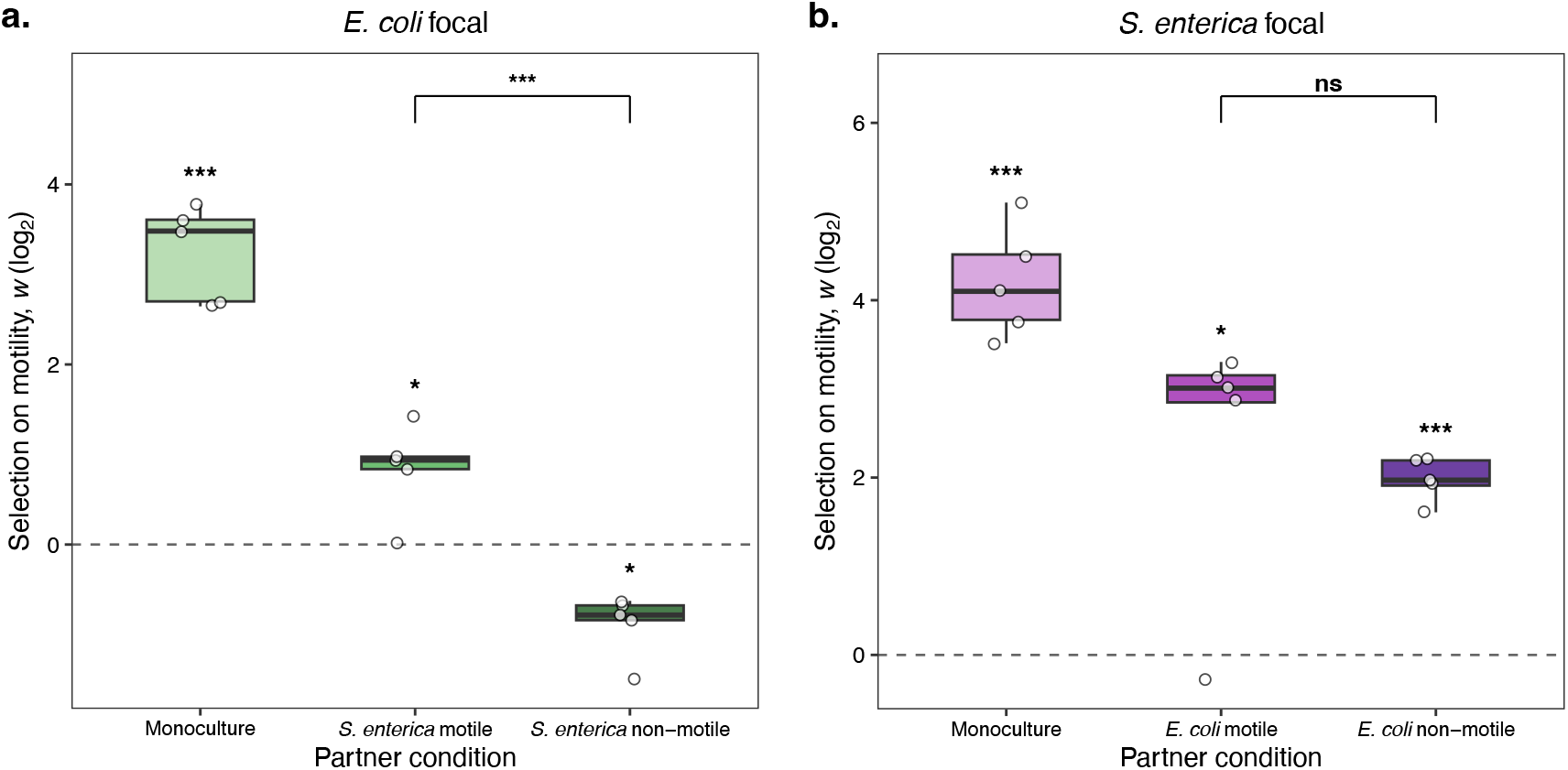
Context dependent selection for motility in spatially structured environments. In panel (a), when *E. coli* is the focal species motility is selected for (*w >* 0) in monoculture and when cross-feeding partner *S. enterica* is motile (middle barplot) but the direction of selection reversed (*w* < 0) and motility is selected against when cross-feeding from non-motile *S. enterica* (right barplot). In panel (b), when *S. enterica* is the focal species, motility is selected for (*w >* 0) in monoculture (left barplot) when cross-feeding with motile *E. coli* (middle barplot) and when cross-feeding with non-motile *E. coli* (right barplots).

In monoculture with methionine supplemented across the whole habitat, motility was strongly favoured (*w* = 3.24 ± 0.24, *p* < 0.001), showing that motility benefits resource acquisition even without a partner. The reversal in co-culture with a non-motile partner therefore reflects a specific effect of the mutualistic interaction.

### *S. enterica* benefits from motility regardless of partner motility

In contrast to *E. coli*, motility was always favoured in *S. enterica* regardless of partner motility status (Figure 2b). With a motile *E. coli* partner, motile *S. enterica* increased in frequency (*w* = 2.41 ± 0.67, t = 3.60, *p* = 0.043, *n* = 5) qualitatively similar in direction to the case where *E. coli* was the focal species in the presence of its motile partner (Figure S2b). However, motility was also favoured in *S. enterica* when the partner was non-motile (*w* = 1.98±0.11, t = 18.26, *p* < 0.001, *n* = 5). Further, we found that selection strength did not differ significantly between partner conditions (Tukey HSD: Δ*w* = 0.436, *p* = 0.7532). As with *E. coli*, monoculture experiments with supplemented acetate showed strong selection for motility (*w* = 4.20 ± 0.28, *t* = 15.02, *p* < 0.001, *n* = 5). The asymmetry between species is therefore not explained by baseline differences in monoculture selection. Rather, we find in our experiments that the two mutualistic partners experience qualitatively different selection regimes in co-culture, namely, that *E. coli* motility selection is partner-dependent, while *S. enterica* selection is not.

### Loss of spatial benefit, not increased motility cost, drives reversal in motility selection in *E*.*coli*

Whether motility is favoured depends not only on the benefit of increased resource access but on whether that benefit offsets the cost of maintaining a motility apparatus. To measure that cost independently, we competed the same strains in well-mixed liquid culture (Figure 3, see also Figure S3). Non-motile strains were favoured in five of six treatments, indicating a generic cost of motility with the exception being *E. coli* with a motile partner. Although not statistically significant, the direction of selection trended in the same way (*w* = −0.91 ± 0.36, *p* = 0.063).

**Figure 3:**
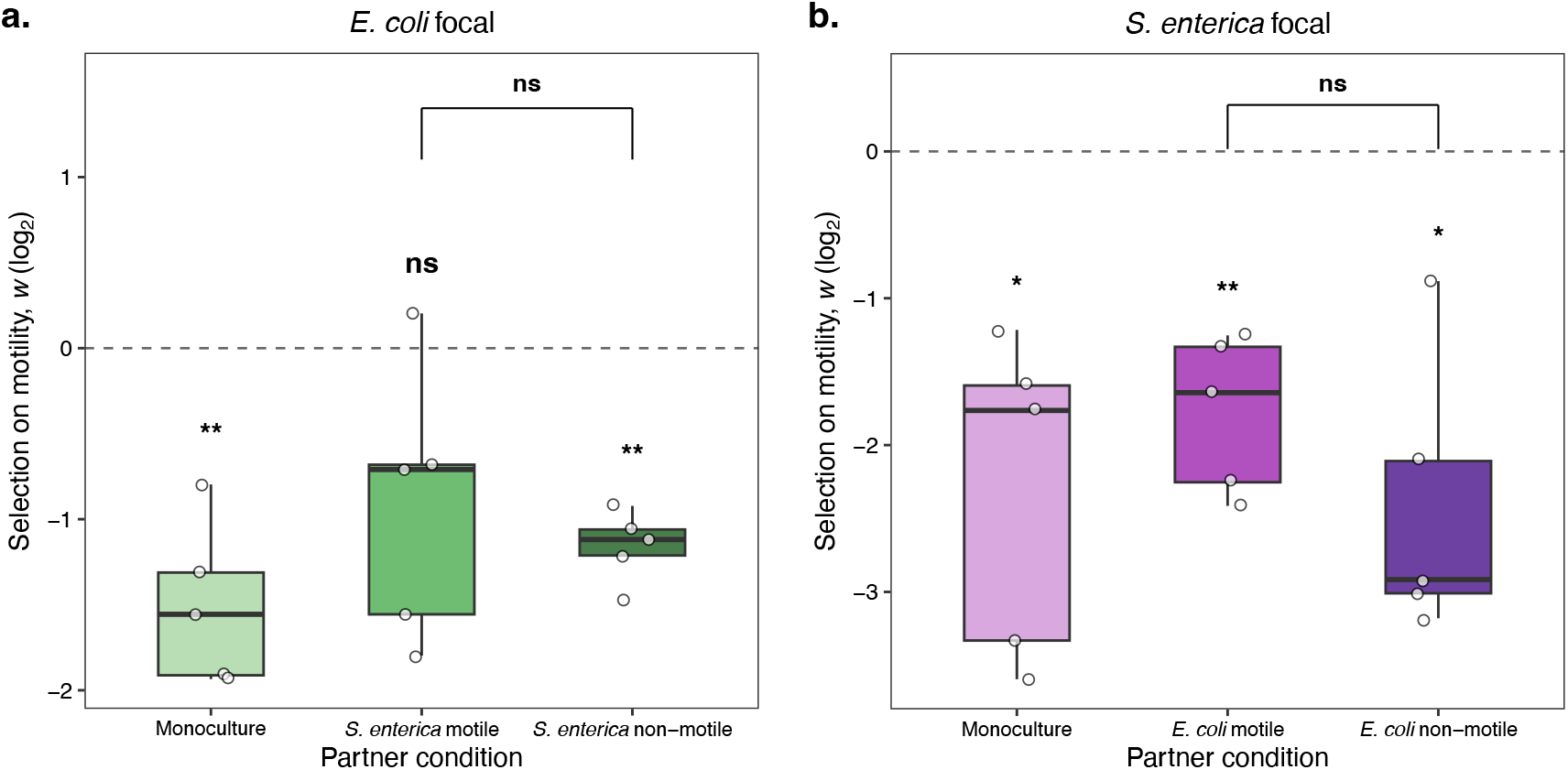
Competition in non-spatial environment almost always selects against motility. The fitness effect of motility was measured in shaking liquid media with *E. coli* (panel A) or *S. enterica* (panel B) as the focal species. For each focal species the fitness of motility was determined in monoculture (left barplot), when cross-feeding from a motile partner (middle barplot), and when cross-feeding from a non-motile partner (right barplot). Motility was strongly selected against in five out of six cases.

**Figure 4:**
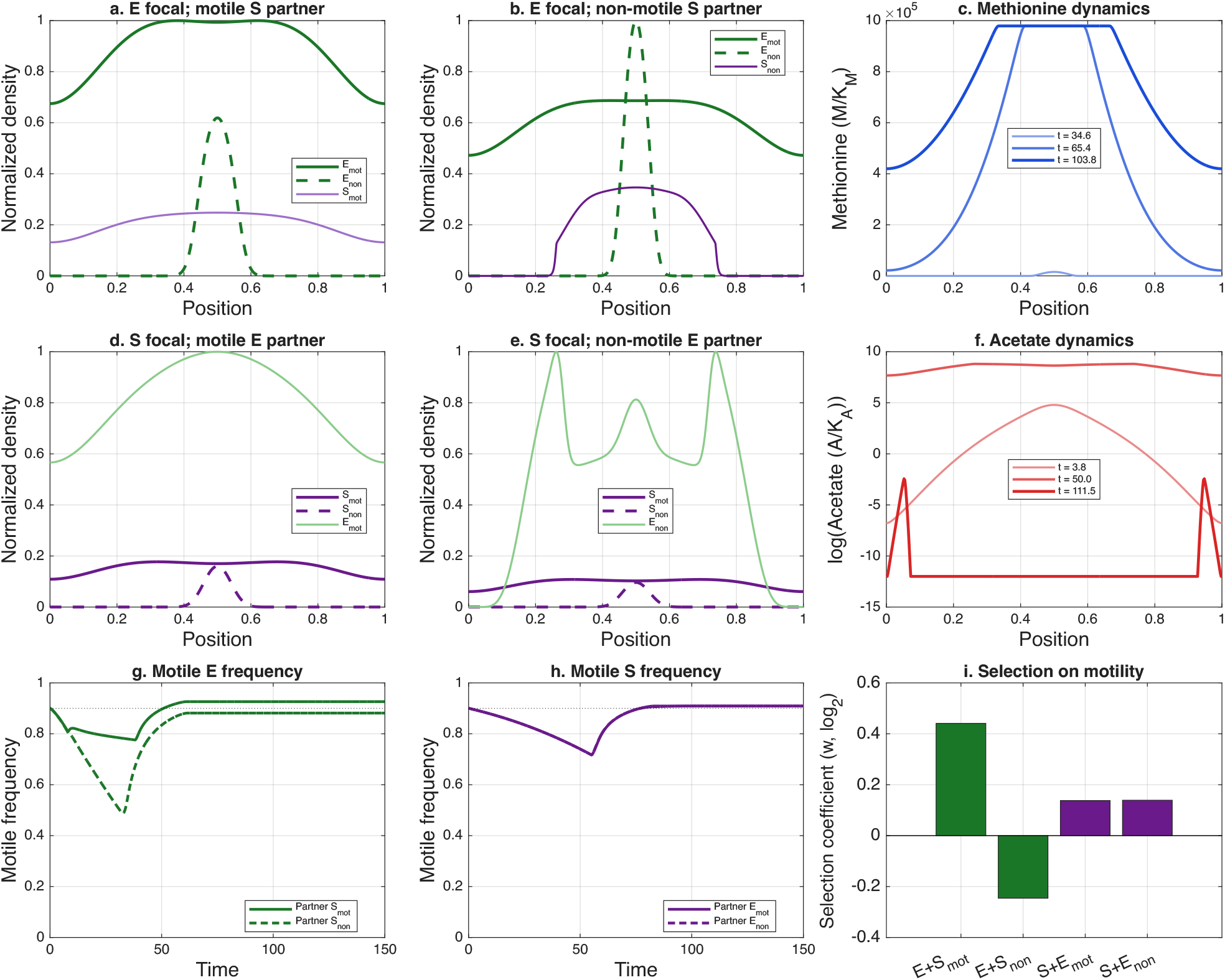
Spatial, resource explicit model qualitatively captures experimental dynamics. Panels (a) and (b) show the endpoint spatial distributions of focal species *E. coli* (‘E’) in the presence of a motile or non-motile *S. enterica* (‘S’) partner respectively. Panel (c) shows spatio-temporal evolution of resource concentration of methionine (blue) across space which are necessary for E growth (where partner is non-motile). Panels (d) and (e) show endpoint spatial distributions of competing focal motile and non-motile strains of *S. enterica* in the presence of motile and non-motile *E. coli* partner respectively. Panel (f) shows the spatio-temporal evolution of acetate dynamics (in log scale; partner is non-motile). In panels (a), (b), (d), (e), density at each point in space is normalised relative the maximal density across both species and spatial domain. Panels (g) and (h) describe the frequency of the motile strain (*E. coli* and *S. enterica* respectively) over time relative to its initial frequency in simulations (0.9). Panel (i) calculates the selection coefficient (w) for each of the four treatments of the simulation and shows qualitatively similar outcomes to experimental selection for motility. Simulations used dimensionless parameters: Da = 9.62 × 10^−4^, *β*_*E*_ = *β*_*S*_ = 0.10, *β*_*p*_ = 0.001, *α* = 0.23, *c* = 0.1, *κ*_*L*_ = 10^−6^, *ρ*_*M*_ = 221, *ρ*_*A*_ = 0.21, *ϕ*_*L*_ = 5.3 × 10^−7^, *ϕ*_*A*_ = 1.7 × 10^3^, *ϕ*_*M*_ = 1.13 × 10^−2^ with characteristic scales *L*_0_ = 4.0 cm, *τ* = 1.54 hr, *E*_0_ = *S*_0_ = 10^6^ cells.

Expressed per generation, this cost was −0.125 ± 0.018, −0.076 ± 0.030 and −0.100 ± 0.007 for *E. coli* in monoculture, with a motile partner and with a non-motile partner respectively, and −0.190 ± 0.042, −0.188 ± 0.026 and −0.269 ± 0.052 for the corresponding *S. enterica* treatments. This estimated per generation cost did not differ between partner conditions in either species (*E. coli*: Δ*s* = 0.024, *p* = 0.47; *S. enterica*: Δ*s* = 0.081, *p* = 0.22), indicating that it was likely generic property of carrying a motility apparatus rather than a feature of the interaction and the specific motility statuses of the partners.

Subtracting this cost from the selection measured on agar (Eq. 2) allowed isolation of the spatial benefit of motility (Table 1). In five of six treatments the benefit was large and significantly positive, from +0.154 ± 0.037 to +0.434 ± 0.045 per generation. The exception was *E. coli* with a non-motile partner, for which the benefit was +0.021 ± 0.016 and not significantly different from zero (*p* = 0.23). The ratio of benefit to cost lay between 1.5 and 2.9 in the five treatments where motility was favoured, but was 0.21 for *E. coli* with a non-motile partner. Thus, we concluded that selection against motility arose not because motility becomes disadvantageous in space or because its cost is prohibitively large, but because the spatial benefit that offsets that cost in every other treatment is absent.

**Table 1:**
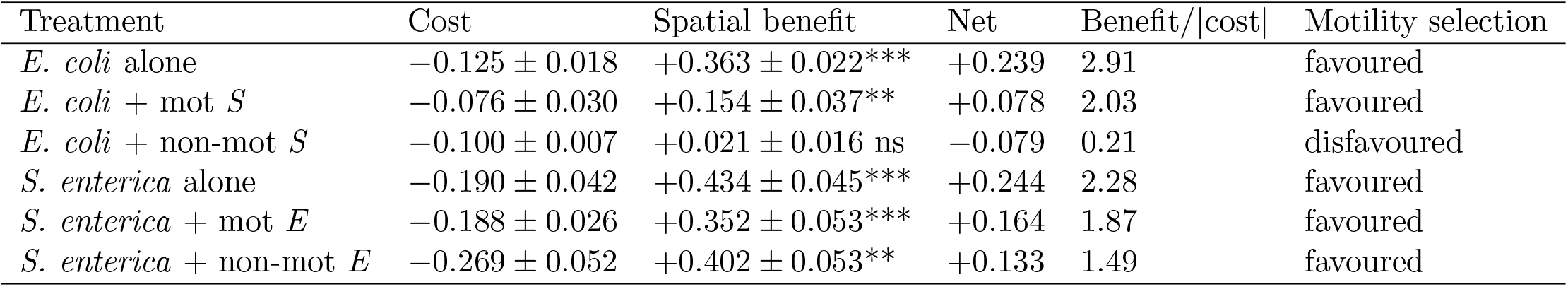
Decomposition of selection on motility into the metabolic cost (measured in well-mixed liquid culture) and the spatial benefit (the additional selection observed on swimming agar). All values are per generation, in log_2_ units, mean ± s.e.m., *n* = 5. The net value is measured directly by the agar assay and equals motility cost plus spatial benefit with only the mean value shown as the difference between benefit and cost (ns:not significant, *:*p* < 0.05, **: *p* < 0.01, ***: *p* < 0.001).

### Metabolite dynamics and growth rate asymmetry explain species-specific selection in a spatially explicit, consumer-resource model

To identify the mechanisms underlying these patterns we simulated the four treatments using the model described above (Figure 1b; Figures S5 and S8). Using empirically derived parameter ranges (Table S1), the model qualitatively recapitulated all four experimental outcomes (Figures **??**i and S4). Methionine accumulated over the course of a simulation (*ρ*_*M*_ = 221, see Figure **??**c) whereas acetate was produced and rapidly consumed, remaining scarce throughout (*ρ*_*A*_ = 0.21, Figure **??**f). This asymmetry in metabolite dynamics is central to the species-level difference in motility selection. To determine more generally how these patterns of motility selection hold and to identify the mechanisms driving them, we systematically varied key dimensionless parameters (described in Methods) and mapped selection on motility across parameter space (Figure 5). Three broad rules emerge from our analysis encapsulating the dynamics in Figures **??** and 5.

**Figure 5:**
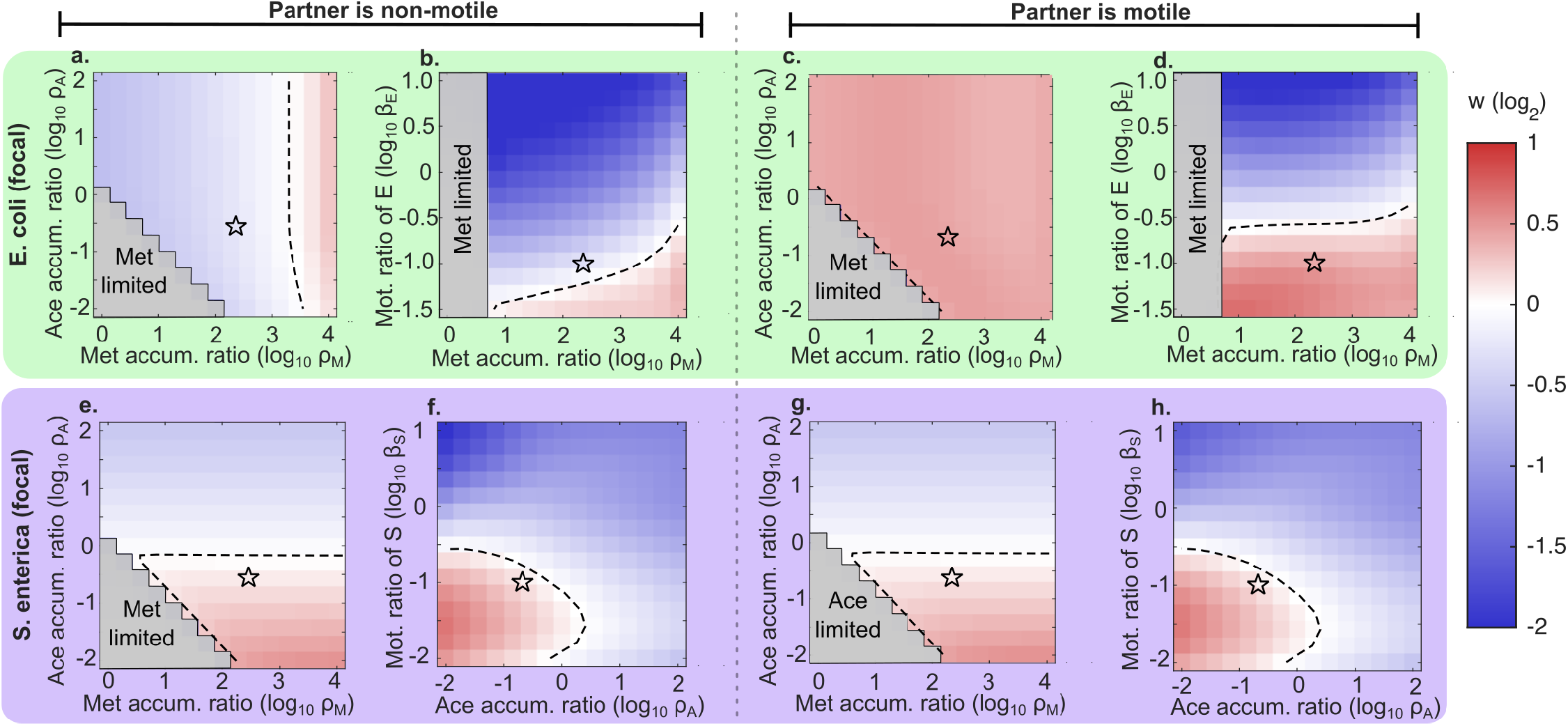
Phase diagram describing the joint effects of metabolite dynamics and motility rates in shaping selection across parameter space. Panels are organised by focal species (top row: *E. coli* ; bottom row: *S. enterica*) and partner motility status (left half shows non motile partner; right half shows motile partner cases). Colour indicates the selection coefficient w, with red denoting selection for motility (*w >* 0), blue denoting selection against motility (*w* < 0), and white indicates neutrality. Stars mark the experimental parameter values (Table S2). Grey shaded regions indicate parameter combinations where a cross-fed metabolite (methionine or acetate) depletes before lactose, placing the system outside the lactose limited regime relevant to our experimental conditions. Panels (a), (c), (e), and (g) vary the methionine accumulation ratio (*ρ*_*M*_ = *λ*_*M*_ */γ*_*M*_, from 10^0^ to 10^4^) against the acetate accumulation ratio (*ρ*_*A*_ = *λ*_*A*_*/γ*_*A*_, from 10^−2^ to 10^2^), with all other parameters held at baseline values. Panels (b) and (d) vary the motility ratio of *E. coli* (*β*_*E*_ = *D*_*E*_*/D*_*M*_, ranging from 10^−1.5^ to 10) against *ρ*_*M*_, with *ρ*_*A*_ and all other parameters at baseline. Panels (f) and (h) vary the motility ratio of *S. enterica* (*β*_*S*_ = *D*_*S*_*/D*_*M*_, from 10^−2^ to 10) against *ρ*_*A*_, with all other parameters at baseline. In each case, the *β* panel pairs a focal species’ dispersal rate with the accumulation ratio of the metabolite it receives from its partner. All simulations were run to steady state using the dimensionless model described in the SI. Constant parameters were held at baseline values: *Da* = 9.62 × 10^−4^, *β*_*E*_ = *β*_*S*_ = 0.10, *β*_*p*_ = 0.001, *α* = 0.23,*c* = 0.1, *κ*_*L*_ = 10^−6^, *ρ*_*M*_ = 221, *ρ*_*A*_ = 0.21, *ϕ*_*L*_ = 5.3 × 10^−7^, *ϕ*_*A*_ = 1.7 × 10^3^, *ϕ*_*M*_ = 1.13 × 10^−2^. Phase diagrams used a 15 × 15 grid. Contour lines depict the zero fitness boundary.

#### Metabolite scarcity creates exploitable gradients

Increasing *ρ*_*M*_ shifted *E. coli* towards selection for motility, while increasing *ρ*_*A*_ shifted *S. enterica* against it (Figure 5a,c,e,g). Both follow from the accumulation ratio of the metabolite each species receives. When that metabolite is scarce relative to demand (*ρ*_*A*_ < 1, as acetate is for *S. enterica*) it is consumed almost as fast as it is produced, and the resulting profile is not a peak at the source but a depletion zone around it. The resource concentration is lowest where consumers are densest and highest ahead of the advancing front, where the metabolite has diffused but no consumers have arrived (Figure **??**f). Because metabolites diffuse an order of magnitude faster than cells (*β* ≈ 0.1), this unexploited reservoir always lies ahead of the front. Motile cells gain not by moving toward the producer but by moving out of the region their own population has exhausted. Conversely, when the metabolite accumulates (*ρ*_*M*_ *ρ*_*A*_ > 1, as for methionine here) it saturates the Monod term wherever consumers are present (*f*_*M*_ → 1; Figure **??**c). Methionine still varies in space, but growth does not, so no exploitable fitness gradient remains. Therefore, we conclude that what selection requires is a gradient not in resource concentration but in the growth rate the resource supports.

#### Co-limitation on multiple resources provides a fallback gradient determining selection

If an exploitable metabolite gradient were the only driver, both species should be disfavoured from motility whenever their cross-fed metabolite is abundant. Instead, at high *ρ*_*M*_ *E. coli* motility is favoured (Figure 5a,c) while at high *ρ*_*A*_ *S. enterica* motility is disfavoured (Figure 5e,g). The difference lies in the number of growth-limiting resources for each species. *E. coli* depends on both lactose and methionine (*r*_*E*_ = *f*_*L*_ *· f*_*M*_ ), so when methionine saturates, growth reduces to *r*_*E*_ ≈ *f*_*L*_, a single-resource regime. The concentrated population then depletes local lactose faster than diffusion replenishes it, creating a consumption-driven depletion front that motile cells can exploit, exactly as in our monoculture cases (where motility is always selected). However, *S. enterica* depends on acetate alone (*r*_*S*_ = *f*_*A*_), so when acetate saturates (*ρ*_*A*_ ≫ 1; numerically explored in Figure 5 and 3 orders of magnitude higher than the experimental value), growth becomes spatially uniform, no alternative gradient exists, and the cost *c* is paid at every timestep. Thus, motility is selected against, in this artificial setting (see Figure S11 for a detailed exploration). To verify this, we simulated a “synthetic” *E. coli* dependent on a single resource (Figure S12). In agreement with our expectation, motility is then favoured regardless of partner status (Figure S12), showing that co-limitation is what makes *E. coli* selection partner-dependent via creation of a spatial gradient.

#### Partner growth and motility set the spatial structure of production

Partner motility expands the region of parameter space favouring motility in *E*.*coli* while it does not particularly change the dynamics of selection in *S. enterica* (Figure 5a versus c, e versus g). This asymmetry arises because for *E. coli*, its motile *S. enterica* partner distributes metabolite production across the spatial domain whereas a non-motile one concentrates it near the inoculation point which creates a ‘partner-dependent’ selection on motility. A non-motile partner can nonetheless generate a distributed source if it grows fast enough to expand by growth alone (in the case where *S. enterica* is the focal species). By interchanging *ρ* between species, where methionine’s accumulation ratio is < 1 and acetate’s is > 1, we can confirm this behaviour (Figure S11). For *E. coli*, partner structure matters because co-limitation requires lactose and methionine at the same location. A motile *S. enterica* partner, on the other hand, distributes methionine so that motile *E. coli* at the periphery meets both fresh lactose and sufficient methionine (Figure **??**a and i). A non-motile partner *S. enterica*, however, remains confined near the centre because it grows slowly (*α* = (*µ*_*S*_*/µ*_*E*_) = 0.23), so methionine stays localised and dispersing *E. coli* encounter lactose without it, reversing selection (Figure **??**b and i). For *S. enterica* the acetate producer grows fast, so even non-motile *E. coli* expands substantially and creates an acetate source that moves outward. Motile *S. enterica* track this front, which is why motility is favoured regardless of partner status (Figure **??**d,e). Growth rate asymmetry (*α* = 0.23) therefore plays a decisive role as a fast-growing partner generates a moving source of benefit even without motility while a slow-growing one remains a static source. Further, we find that growth rate ratio *α* and cost *c* act as a threshold (Figures and S6), with low costs favouring motile strains regardless of *α*.

#### Motility benefits are bounded by the spatial distribution of partner-derived metabolites

Varying each species’ dispersal rate against the accumulation ratio of the metabolite it receives reveals a ceiling on beneficial motility (Figure 5b,d,f,h). Because *β* compares bacterial dispersal to metabolite diffusion, cells with high *β* outrun the region in which the cross-fed metabolites essential to them are present. As *β* → 1 individuals overshoot the location of the metabolite they depend on. In every treatment the zero contour lies below *β* = 1 i.e. (log_10_*β*) = 0, at *β* ≤ 0.3, since cells must remain well inside the zone of metabolite presence rather than at its edge while paying the cost of motility throughout.

The position of this “ceiling” on motility differs between species in a way that also follows from co-limitation. For *E. coli* it rises with *ρ*_*M*_ i.e. a larger methionine pool extends the region of motility benefit, so faster dispersal remains profitable, and a motile partner increases this region further by distributing production across space (Figure 5d versus b). The favourable region is therefore present across all values of *ρ*_*M*_ . In contrast, for *S. enterica* the favourable region for motility is instead bounded at a maximal *β*_*S*_ and to the right by acetate saturation (large *ρ*_*A*_; Figure 5f,h), because no second resource provides an alternative gradient once acetate saturates.

These results resolve the two hypotheses posited in Figure 1c. *E. coli* follows H1, in which matching the partner’s motility status is favoured, while *S. enterica* follows H2, in which motility is favoured irrespective of the partner. Neither hypothesis holds for the community as a whole, and the two partners in the same interaction follow different predictions due to the mechanisms proposed here.

## Discussion

Natural microbial communities contain diverse cross-feeding interactions embedded within networks of cooperation, competition, and exploitation [15, 37, 38]. In this paper, we ask a simple question: how and when does motility get selected for in cross-feeding microbial mutualisms? Our results show that cross-feeding mutualisms generate asymmetric, context-dependent selection on motility between partners. This asymmetry is driven by three interacting features of the metabolic interaction namely the scarcity of the cross-fed metabolite, the number of growth-limiting resources, and the partner’s spatial behaviour. These rules provide a predictive framework for motility evolution in mutualistic communities.

### Motility selection depends on metabolite dynamics

Our findings extend existing theory on motility evolution in mutualisms across micro and macro-scale ecologies. Previous models predicted that dispersal should be disfavoured in obligate mutualisms because individuals that move away from their partners lose access to essential benefits [25–27]. Our results show that this logic does not always hold for cross-feeding mutualisms, where benefits diffuse through the environment. When metabolites can spread, the relevant question is not how proximal the partner is, but where metabolites are abundant. When the cross-fed metabolite accumulates (for e.g. *ρ*_*M*_ > 1 in our system), it saturates the domain regardless of the consumer’s position, eliminating the need for motility. Recent experiments with *Pseudomonas* showed that when *P. putida* evolved in co-culture with a cross-feeding partner, functional motility was maintained and even enhanced, whereas the same populations rapidly lost motility when the identical resource was supplied externally and uniformly [39]. This parallel between biogenic (partner-produced and spatially structured) and non-biogenic (externally supplied and uniform) resources maps directly onto our modelling framework. Essentially, motility selection is determined by the accumulation ratio of the metabolite the focal species itself receives, which sets whether that metabolite is drawn down as fast as it arrives or builds to saturation. Biogenic supply resembles the low-*ρ* regime, in which gradients persist and motility is favoured whereas uniform external supply resembles the saturating or high-*ρ* regime, in which it is not.

Our model thus provides a mechanistic basis for the context-dependent maintenance of motility observed across cross-feeding systems. Thus, the distinction between localised and diffusible benefits may apply broadly with any mutualism mediated by an environmentally exchanged benefit (e.g. siderophores, signalling molecules, amino acids) and should exhibit dynamics closer to our cross-feeding case than to mutualism models assuming physical proximity of partners. Indeed, while past research suggests that environmentally mediated metabolite exchanges often only arise at the scale of a few cell-lengths, a systematic characterisation of spatial metabolite profiles in ecological communities remains an open problem [23].

Decomposing selection into its cost and spatial components clarifies what changes when a partner cannot move. The metabolic cost of motility was invariant across partner conditions, so the reversal we observe in *E. coli* is not explained by the cost of flagella becoming heavier in the presence of a non-motile partner. Instead the spatial benefit reduces in these treatments. A non-motile, slow-growing *S. enterica* partner (*α* = 0.23) confines methionine production close to the inoculation point, so motile *E. coli* that disperse outward encounter lactose without the methionine they also require. Because *E. coli* is co-limited, movement into such regions yields no growth, and motility ceases to provide any expected benefit. The cost, which is paid regardless, then determines the selection outcome. Thus, the benefit shrinks relative to the cost to the point where it becomes almost undetectable resulting in selection against motility.

### Variation in motility as a mechanism of strain coexistence

Finer probing of microbial communities, with advances in metagenomics and other sequencing techniques, highlights extensive fine-grained diversity found in these systems [40–42]. Indeed, several strains within a species are known to coexist with one another despite having likely high overlaps in their ecological niches [43]. Several mechanisms, such as differences in metabolic activity, physiological trade-offs or susceptibility to phage predation have been invoked but these often ignore the role of spatial structure in shaping ecological dynamics of a community [44–46]. Natural environments such as the human gut, skin or nasal passages are all also spatially structured thereby shaping the dynamics of community assembly [40, 41, 47]. Indeed, Al-Tameemi and Rodríguez-Verdugo [48] find that when *P. putida* diversifies into motile and non-motile morphotypes, the non-motile type sweeps to fixation in monoculture but both morphotypes coexist stably for approximately 150 generations when the cross-feeding partner is present. The partner thus stabilises motility polymorphism, consistent with our finding that partner-generated spatial heterogeneity creates the conditions for frequency-dependent coexistence (Figure S7). Negative frequency dependence of this form can also maintain polymorphism within populations [49–51], suggesting that motile and non-motile strains may stably coexist rather than one replacing the other. This is consistent with empirical observations of motility variation within natural bacterial populations and with recent demonstrations that growth-motility tradeoffs can sustain strain coexistence in structured habitats [16, 52]. More broadly, at the single-cell scale, motility enhances spatial inter-mixing and disrupts clonal clustering in cross-feeding consortia, increasing both individual growth rates and community productivity [28].

### Implications for community assembly

If motility selection depends on partner motility status, as we have shown for *E. coli*, then the community’s motility composition feeds back on to selection. A community dominated by motile partners would favour motility in *E. coli* whereas a community of non-motile partners would disfavour it. This feedback could generate alternative community states or constrain community assembly trajectories, recapitulating priority effects observed in spatially structured microbial communities [53,54]. Motility benefits in cross-feeding systems could also extend beyond spatially structured surface as evidenced in turbulent aquatic environments where swimming enhances partner encounters and adhesion in interkingdom mutualisms [29]. More broadly, our results predict that communities dominated by scarce, rapidly consumed cross-fed metabolites (*ρ* < 1) will consistently favour motile phenotypes, while communities where exchanged metabolites accumulate (*ρ >* 1) will exhibit weaker or partner-dependent selection. Testing these predictions would require characterising both metabolite dynamics and diffusibility across diverse interaction types, an increasingly feasible goal with modern spatially resolved metagenomics and metatranscriptomics [55, 56].

### Extensions and conclusions

We believe several extensions merit investigation. First, our model assumed undirected motility, and while this conservative approach recapitulated experimental results bacteria commonly exhibit chemotaxis toward nutrients and partner-produced metabolites [17, 57]. Chemotaxis should strengthen selection for motility by reducing the cost of misdirected movement, but could also change the conditions under which motility is favoured. If cells can precisely track metabolite gradients, the benefit of motility may persist even when metabolites accumulate globally. Second, our experiments initialised partners at the same location but different spatial configurations could alter selection if they change the effective distance to partner-produced metabolites [30]. Recent theoretical work on mutualist co-invasion suggests that dispersal asymmetries can generate complex coexistence patterns even between obligate partners [58]. Third, extending this framework to facultative mutualists would test how obligacy modulates selection and whether metabolite dynamics dominate regardless of the strength of dependence [59]. Lastly, extending beyond metabolites to other exchanged goods such as vitamins, which are required in trace amounts but expensive to synthesize [60, 61], could test whether our framework generalises to exchanges with different costs and stoichiometries.

In summary, we show that cross-feeding mutualisms generate asymmetric, context-dependent selection on motility between partners. This selection depends on metabolite accumulation dynamics, partner motility status, and growth rate asymmetries which are factors not captured by existing theory developed for localised mutualisms. More broadly, our results illustrate that the evolution of spatial behaviours like motility cannot be understood without considering how species interactions restructure the resource landscape. Thus, in spatially structured communities, the traits of the interacting partners can significantly impact the direction of evolution.

## Acknowledgements

NV would like to thank members of the Harcombe lab particularly Sen Xiong and Ave Bisesi for patiently answering basic queries about microbial experimental protocols. NV also thanks Aarcha Thadi for workshopping parts of Figure 1. This manuscript is work supported by the National Science Foundation under Grant No. DEB-2109965.

## Supplementary Information

Naven Narayanan Venkatanarayanan, Jonathan N.V. Martinson, Allison K. Shaw, William R. Harcombe

### S1 Complete mechanistic model of cross-feeding mutualism

We model the spatial dynamics of a cross-feeding mutualism between *E. coli* and *S. enterica* using a system of reaction-diffusion equations. *E. coli* consumes lactose and produces acetate as a byproduct, while *S. enterica* consumes acetate and produces methionine. *E. coli* requires methionine for growth, establishing an obligate cross-feeding interaction.

Each species exists as two competing strains: a motile strain (subscript “mot”) that actively disperses but pays a growth rate cost, and a non-motile strain (subscript “non”) that disperses only through passive diffusion but grows at the maximum rate. The state variables are *E*_mot_, *E*_non_, *S*_mot_, *S*_non_ (cells cm^−1^) and the resource concentrations *L, A, M* (g L^−1^).

The population dynamics are governed by:

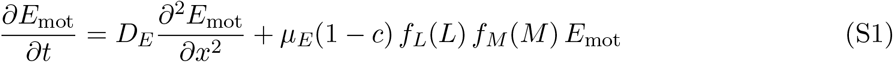

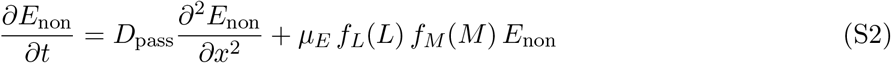

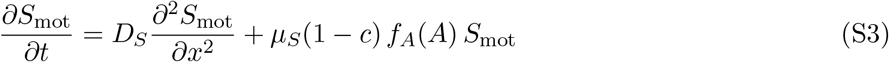

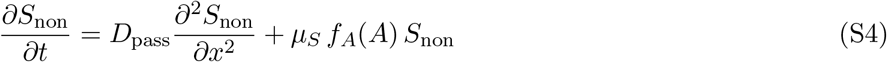

where *D*_*E*_ and *D*_*S*_ are the effective diffusion coefficients for motile *E. coli* and *S. enterica* respectively, *D*_pass_ is the passive diffusion coefficient for non-motile strains, *µ*_*E*_ and *µ*_*S*_ are maximum growth rates, and *c* is the fractional cost of motility.

Growth is limited by nutrient availability through Monod kinetics:

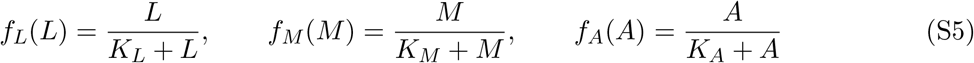

where *K*_*L*_, *K*_*M*_, and *K*_*A*_ are half-saturation constants.

Metabolic flux is proportional to the realised growth rate of each strain. A motile cell divides at a rate reduced by (1 − *c*) and therefore consumes and produces at a correspondingly reduced rate, so the per cell consumption and production terms are weighed by

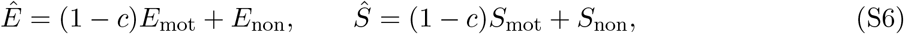

rather than by the raw totals. Note that using the unweighted totals would make a motile cell consume the same lactose per unit time while dividing more slowly, which is equivalent to imposing a yield penalty of *c/*(1 − *c*) on top of the intended rate cost. Because the strains differ only in dispersal and in *c*, that second penalty would confound the cost of motility with a cost of inefficiency, and it would feed back on the resource landscape that generates the selection we measure. The weighted form isolates *c* as a pure growth rate cost.

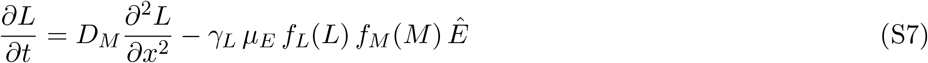

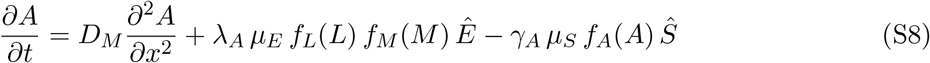

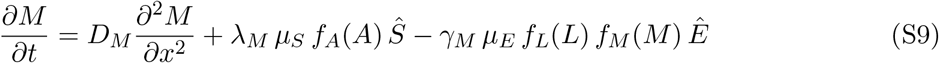

Here *D*_*M*_ is the diffusion coefficient for metabolites (assumed equal for all small molecules), *λ*_*A*_ and *λ*_*M*_ are production yields (g metabolite per cell division), and *γ*_*L*_, *γ*_*A*_, and *γ*_*M*_ are consumption yields (g metabolite per cell division). This formulation ensures that non-growing cells neither consume nor produce metabolites and resource flux is strictly proportional to growth rate. The system is solved on *x* ∈ [0, *L*] with no-flux boundary conditions, *∂u/∂x* = 0 at *x* = 0 and *x* = *L*, for all state variables.

### S2 Non-dimensionalisation of spatially explicit cross-feeding model

- We introduce characteristic scales to non-dimensionalise the model:
- Length scale: *L*_0_ = ℒ (domain length)
- Time scale: *τ* = 1*/µ*_*E*_ (inverse of *E. coli* growth rate)
- *E. coli* density scale: *E*_0_ (initial total cell number)
- *S. enterica* density scale: *S*_0_ (initial total cell number)
- Lactose scale: *L*_scale_ = *L*_init_ (initial lactose concentration)
- Acetate scale: *A*_scale_ = *K*_*A*_ (half-saturation constant)
- Methionine scale: *M*_scale_ = *K*_*M*_ (half-saturation constant) We define dimensionless variables (denoted with tildes):

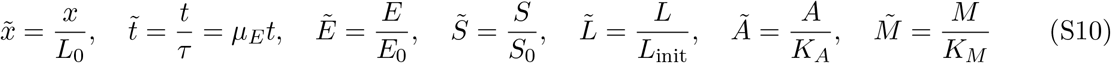

Substituting into the governing equations and dropping tildes for clarity, the dimensionless population dynamics become:

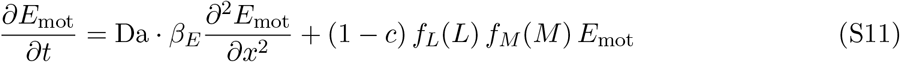

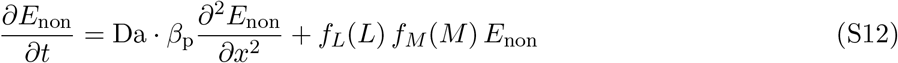

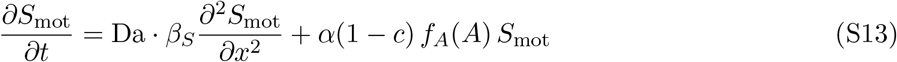

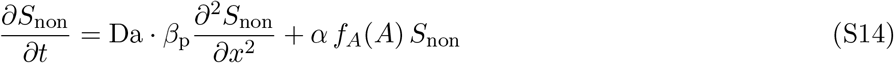

The dimensionless Monod functions are:

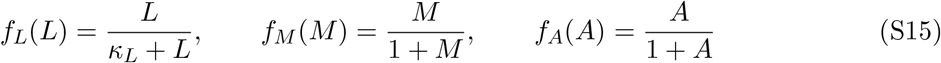

where *κ*_*L*_ = *K*_*L*_*/L*_init_. Note that *f*_*M*_ and *f*_*A*_ simplify because we scaled *M* by *K*_*M*_ and *A* by *K*_*A*_. The dimensionless resource dynamics, with consumption and production coupled to growth, are:

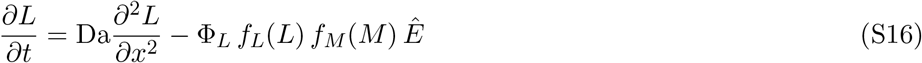

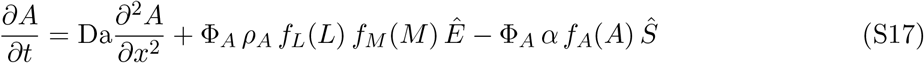

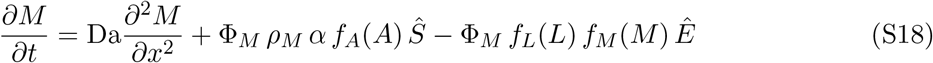

Note that *α* appears in the *Ŝ* terms because *S. enterica*’s growth rate is *µ*_*S*_ = *αµ*_*E*_, and this growth rate multiplies the consumption/production yields in the dimensional equations. When we non-dimensionalise time by *τ* = 1*/µ*_*E*_, the factor *µ*_*E*_*τ* = 1 cancels for *E. coli* terms, while *µ*_*S*_*τ* = *α* remains for *S. enterica* terms.

The dimensional parameters are defined in Table S1, and the corresponding dimensionless parameter values is given in Table S2.

### S3 Numerical implementation

The system was solved with a finite difference method on the dimensionless domain *x* ∈ [0, 1], discretised with *N* = 401 uniformly spaced grid points and spacing Δ*x* = 1/(*N* − 1) = 2.5 × 10^−3^. The Laplacian used second-order central differences at interior points, and no-flux boundaries were imposed, giving

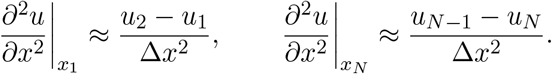

We used an implicit-explicit scheme for solving the coupled differential equations. Diffusion and production terms were advanced explicitly and the resource consumption terms which were stiffest, were advanced implicitly. Because every consumption term is of the Monod form Φ *u/*(*κ* + *u*), we can write it as *Cu* with *C* evaluated at the previous step, giving a linearly implicit update that requires no iteration. For a resource *u* with diffusion coefficient *D*, production *P* and consumption coefficient *C*,

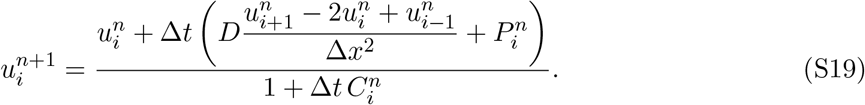

Population densities, whose reaction terms are not stiff, were advanced explicitly. Simulations were monitored for non-finite values and terminated if any were detected.

Both bacterial populations were initialised with a Gaussian centred at *x* = 0.5 of width *σ* = 0.03, normalised to the unit integral. Because *σ* is specified in physical rather than grid units, the inoculum does not change size when the mesh is refined and the so mesh refinement changes only the numerical resolution. In competition experiments, the focal species was initialised at 90% motile and 10% non-motile, and the partner species entirely motile or entirely non-motile according to treatment. Lactose was initialised uniformly at *L* = 1. Acetate and methionine were initialised with the same Gaussian profile, scaled to a peak value of unity in units of their respective half-saturation constants, reflecting production by the localised initial populations. In the supplemented monoculture runs (Figure S4) the cross-fed metabolite the focal species requires was instead supplied uniformly across the spatial domain of the model (mimicking the experiments), and lactose was set to zero in the *S. enterica* monoculture, which cannot use it.

Each simulation was run for a fixed dimensionless window *t* = 150. We use a fixed window rather than a resource-depletion criterion for two reasons. First, treatments differ in how quickly they exhaust the supplied lactose, so a depletion rule would compare selection integrated over different durations and reporting a common window removes that confound. This fixed time window also lets *w* be normalised by the doublings each treatment actually completed. Second, acetate is *S. enterica*’s only resource and is always fully consumed but this happens often after *E. coli* has run out of lactose and stopped producing it, so resources depletion-time criterion will erroneously report the selection outcome before steady state. In all our simulations, the total population change over the final third of the window is below 0.1% in all four co-culture treatments, and the endpoint population reaches the analytic bound 1 + 1*/*Φ_*L*_ imposed by the supplied lactose. We also made sure that the selection coefficient ‘w’ converged within a bound of 10^−4^ units at the end of the simulation.

### Selection

Selection on motility was quantified exactly as in the experiments,

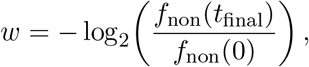

with *f*_non_ the frequency of non-motile cells and populations integrated over space. Positive *w* indicates selection for motility.

**Figure S1:**
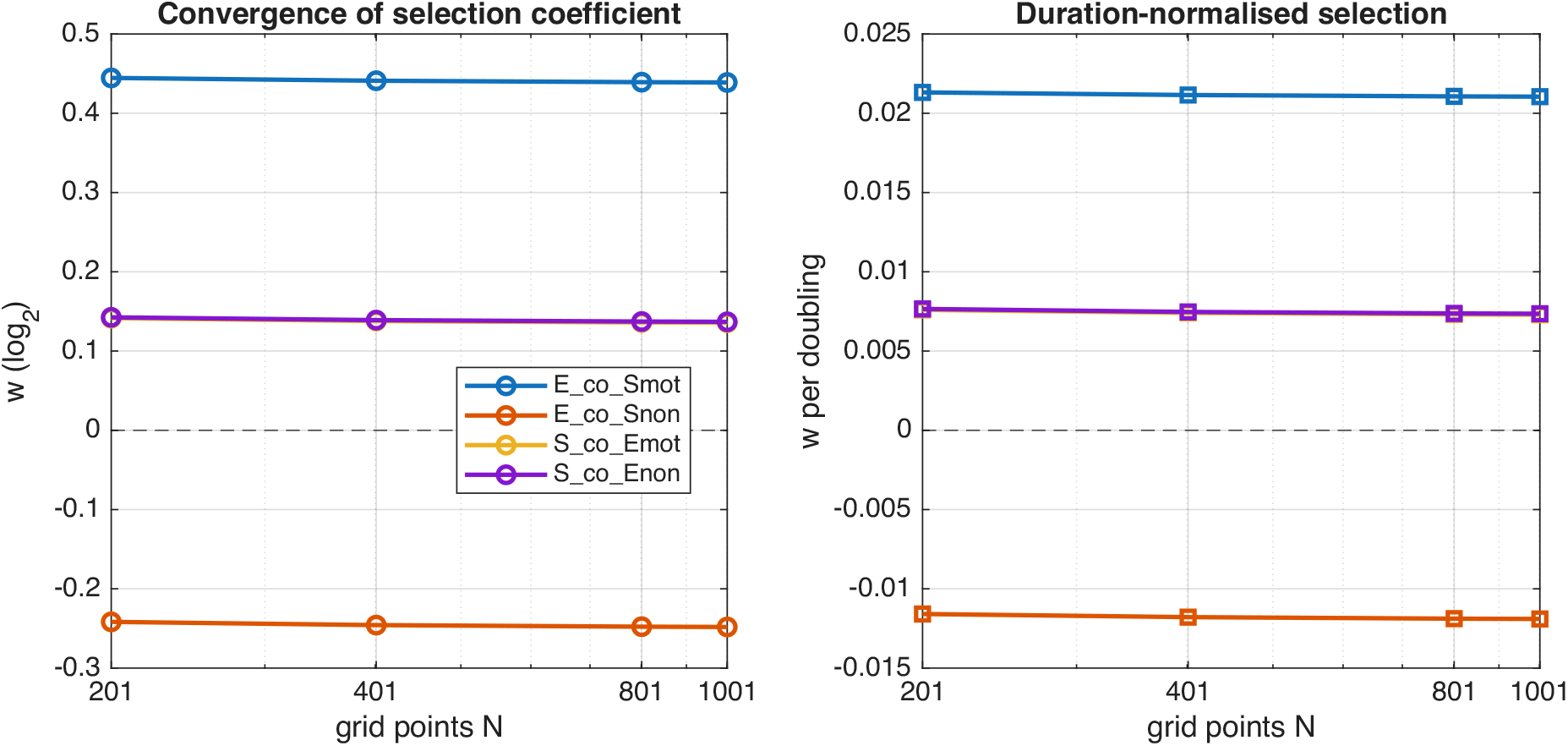
Grid convergence of the selection coefficient. We performed all four co-culture treatments at *N* = 201, 401, 801 and 1001 grid points, holding the inoculum width fixed in physical units so that mesh refinement changes only the numerical resolution, and scaling Δ*t* to satisfy the diffusive stability limit at every resolution. We find that the selection coefficient remains the across all mesh sizes indicating convergence. Note that the purple (S co Enon) and yellow (S co Emot) are overlaid on top of one another. All other parameters are at their baseline values (Table S2). Panel on left shows cumulative selection coefficient *w* (log_2_). Panel on right shows the same quantity normalised by the number of focal-population doublings. Results reported in the main text use *N* = 401.

### Grid convergence of selection estimates

To make sure that our results were not an artefact of the choice of grid size, we verified convergence by repeating all four treatments at N = 201, 401, 801, and 1001 grid points, scaling Δ*t* so that Δ*t/*Δ*x*^2^ remained fixed. Show in FigureS1 is the convergence of the selection coefficient at the end of the simulations (fixed time; left panel) as well as coefficients normalised by number of doublings. All results reported in the main text use *N* = 401. At coarser resolutions the non-motile population spans only one to two grid cells and selection estimates are correspondingly unreliable, particularly for treatments in which *w* is small.

**Table S1:** Dimensional parameters used in the cross-feeding model. All concentrations are in g/L and all yields in g per cell division.

| Parameter | Symbol | Value | Source |
| --- | --- | --- | --- |
| *Diffusion coefficients* |  |  |  |
| Metabolite diffusion | $D_M$ | 0.01 cm<sup>2</sup>/hr | adapted from [16] |
| *E. coli* motile diffusion | $D_E$ | 0.001 cm<sup>2</sup>/hr | adapted from [16, 31] |
| *S. enterica* motile diffusion | $D_S$ | 0.001 cm<sup>2</sup>/hr | adapted from [16, 31] |
| Passive diffusion | $D_{\text{pass}}$ | 10<sup>-5</sup> cm<sup>2</sup>/hr | assumed |
| *Growth rates* |  |  |  |
| *E. coli* max growth rate | $\mu_E$ | 0.65 hr<sup>-1</sup> | [62] |
| *S. enterica* max growth rate | $\mu_S$ | 0.15 hr<sup>-1</sup> | [62] |
| Motility cost | $c$ | 0.1 | assumed |
| *Half-saturation constants* |  |  |  |
| Acetate | $K_A$ | 10<sup>-9</sup> g/L | adapted from [31] |
| Methionine | $K_M$ | 10<sup>-6</sup> g/L | adapted from [31] |
| Lactose | $K_L$ | 10<sup>-6</sup> g/L | adapted from [31] |
| *Production yields* |  |  |  |
| Acetate production | $\lambda_A$ | 3.6 × 10<sup>-13</sup> g/cell | Chacón & Hammarlund |
| Methionine production | $\lambda_M$ | 2.5 × 10<sup>-12</sup> g/cell | effective value; see note |
| *Consumption yields* |  |  |  |
| Acetate consumption | $\gamma_A$ | 1.7 × 10<sup>-12</sup> g/cell | Chacón & Hammarlund |
| Methionine consumption | $\gamma_M$ | 1.13 × 10<sup>-14</sup> g/cell | Chacón & Hammarlund |
| Lactose consumption | $\gamma_L$ | 5.3 × 10<sup>-13</sup> g/cell | Chacón & Hammarlund |
| *Initial conditions* |  |  |  |
| Domain length | $L$ | 4.0 cm | assumed |
| Initial cell number | $N_{\text{init}}$ | 10<sup>6</sup> cells | see note |
| Initial lactose | $L_{\text{init}}$ | 1 g/L | assumed |

### Model Parameters

#### Note on *λ*_*M*_

Direct measurement of methionine production in spent media gave a *λ*_*M*_ ≈ 2.5 × 10^−14^ g per cell division. This value is a lower bound for two reasons. Firstly, the assay is terminated by lactose limitation before methionine production is complete, and it captures only the growth-coupled component of secretion, whereas *S. enterica* likely releases methionine independently of division. Applied directly, it would give *ρ*_*M*_ *ρ*_*A*_ < 1, under which the exchange is not self-sustaining and the co-culture could not grow, contradicting the observed mutualism. We therefore use an effective production rate of 2.5 × 10^−12^ g per cell division, of the same order as the value used by [62], which absorbs growth-independent secretion into the growth-coupled term. Because this parameter is uncertain, we varied *ρ*_*M*_ over four orders of magnitude (Figure 5a and c) and the results reported here generally hold wherever the mutualism is self-sustaining (i.e. *ρ*_*M*_ *ρ*_*A*_ *>* 1).

#### Note on *N*_init_

Simulations were initialised with 10^6^ cells per species as a model normalisation. The experimental inoculum was approximately 10^4^ cells per species but because the model runs to lactose depletion, the endpoint is insensitive to this choice.

**Table S2:** Dimensionless parameters derived from Table S1. All quantities are dimensionless; the reaction volume is absorbed into the concentration scales of Table S3.

| Parameter | Definition | Value | Interpretation |
| --- | --- | --- | --- |
| Da | $D_M/(\mu_E L^2)$ | $9.62 \times 10^{-4}$ | Diffusion vs. growth timescale |
| $\beta_E$ | $D_E/D_M$ | 0.10 | *E. coli* motility ratio |
| $\beta_S$ | $D_S/D_M$ | 0.10 | *S. enterica* motility ratio |
| $\beta_p$ | $D_{\text{pass}}/D_M$ | 0.001 | Passive diffusion ratio |
| $\alpha$ | $\mu_S/\mu_E$ | 0.23 | Growth rate ratio |
| $c$ | — | 0.1 | Motility cost |
| $\kappa_L$ | $K_L/L_{\text{init}}$ | $10^{-6}$ | Lactose saturation |
| $\rho_M$ | $\lambda_M/\gamma_M$ | 221.23 | Methionine accumulation ratio |
| $\rho_A$ | $\lambda_A/\gamma_A$ | 0.212 | Acetate accumulation ratio |
| $\Phi_L$ | $\gamma_L N_{\text{init}}/L_{\text{init}}$ | $5.3 \times 10^{-7}$ | Lactose consumption capacity |
| $\Phi_A$ | $\gamma_A N_{\text{init}}/K_A$ | $1.7 \times 10^3$ | Acetate consumption capacity |
| $\Phi_M$ | $\gamma_M N_{\text{init}}/K_M$ | $1.13 \times 10^{-2}$ | Methionine consumption capacity |

**Table S3:** Characteristic scales used for non-dimensionalisation.

| Scale | Symbol | Value |
| --- | --- | --- |
| Length | $L_0$ | 4.0 cm (domain length) |
| Time | $\tau$ | $1/\mu_E = 1.54$ hr |
| *E. coli* density | $E_0$ | $10^6$ cells |
| *S. enterica* density | $S_0$ | $10^6$ cells |
| Lactose concentration | $L_{\text{scale}}$ | $L_{\text{init}} = 1$ g/L |
| Acetate concentration | $A_{\text{scale}}$ | $K_A = 10^{-9}$ g/L |
| Methionine concentration | $M_{\text{scale}}$ | $K_M = 10^{-6}$ g/L |

### S4 Estimating number of generations for liquid and swimming agar experiments

**Table S4:** Mean focal-population doublings in each assay. We assume both assays are terminated by exhaustion of the same lactose supply, so the number of doublings is set by the lactose yield relative to the inoculum rather than by elapsed time. We find that the 24h liquid assay and a five-day plate assay give comparable generation counts.

| Treatment | Liquid (24h) | Agar (5d) |
| --- | --- | --- |
| *E. coli* alone | 12.1 | 13.5 |
| *E. coli* + motile *S* | 11.9 | 10.7 |
| *E. coli* + non-motile *S* | 11.5 | 11.2 |
| *S. enterica* alone | 12.2 | 17.2 |
| *S. enterica* + motile *E* | 9.5 | 14.6 |
| *S. enterica* + non-motile *E* | 9.1 | 14.9 |

### S5 Supplementary figures

The supplementary figures in this section are divided into sections corresponding to main text figure related to them.

**Figure S2:**
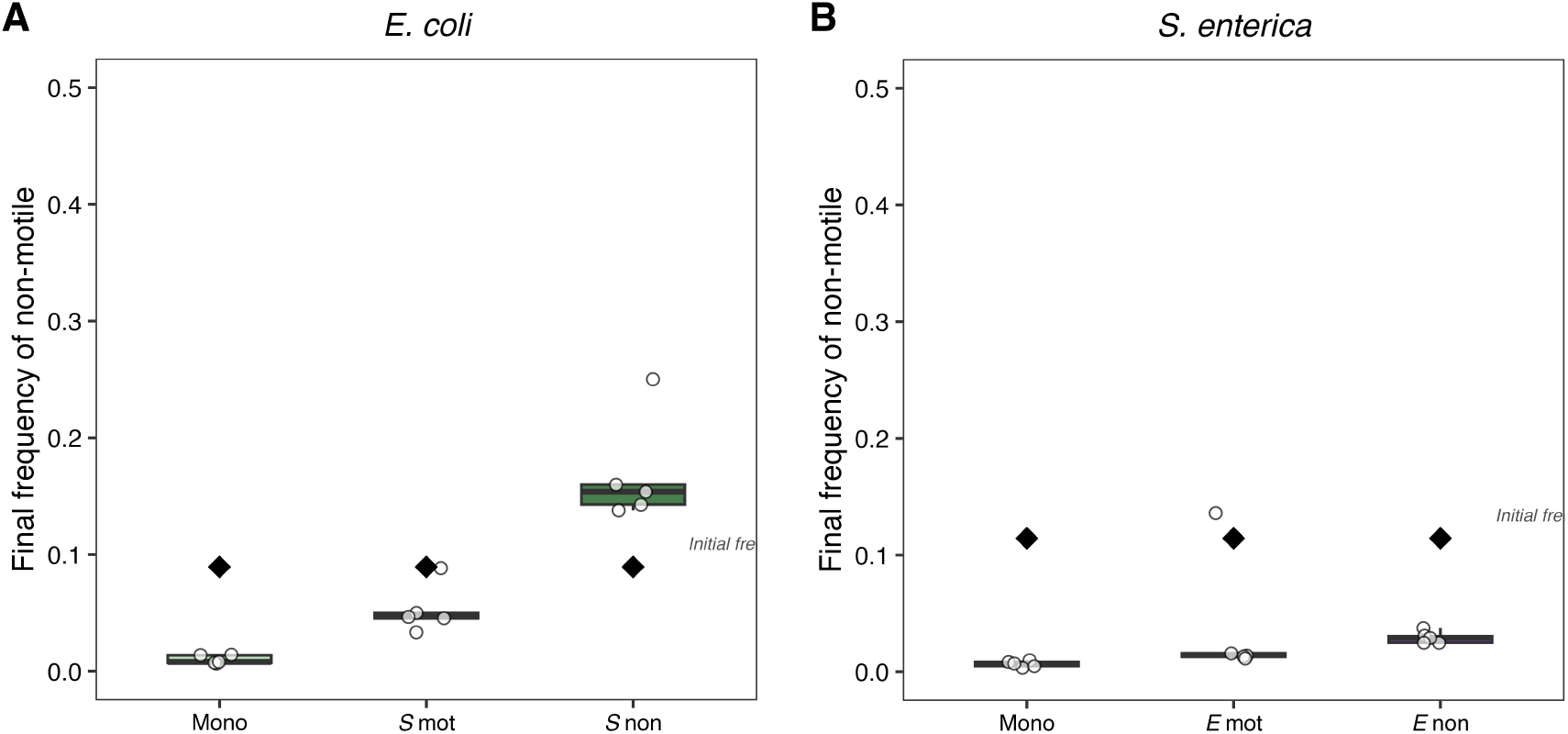
Final frequency of focal species strains in the swimming media experiments. Fitness of motile and non-motile strains calculated in Figure 2 derived from these experiments. Panel (A) shows data for *E. coli* being the focal species and panel (B) shows data for *S. enterica* when it is the focal species

#### S5.1 Supplementary figures associated with Figure 2

Figure shows the change in frequency from initial frequency (black diamond) to final frequency in each of the treatments for the swimming experiments for *E. coli* in panel A and *S. enterica* in panel B.

**Figure S3:**
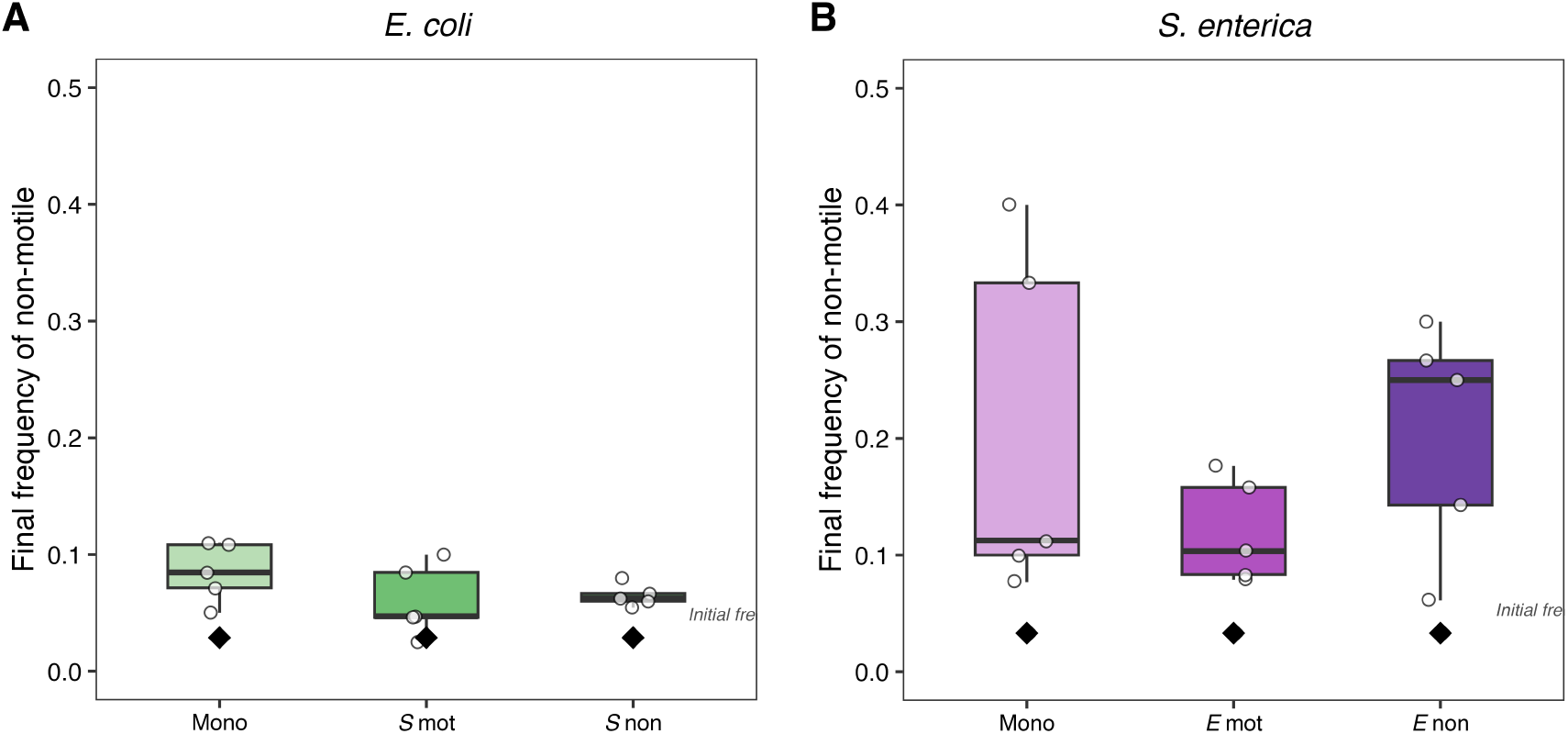
Final frequency of focal species strains in the liquid culture experiments. Fitness of motile and non-motile strains calculated in Figure 2 derived from these experiments. Panel (A) shows data for *E. coli* being the focal species and panel (B) shows data for *S. enterica* when it is the focal species

#### S5.2 Supplementary figures associated with Figure 3

Figure shows the change in frequency from initial frequency (black diamond) to final frequency in each of the treatments for the liquid culture experiments for *E. coli* in panel A and *S. enterica* in panel B.

#### S5.3 Supplementary figures associated with Figure 4

In this section we further show the dynamics of our spatial, resource explicit mathematical model. In particular we present four supplementary results in addition to the ones presented in the main text (Figure **??**) focused on the temporal dynamics of the strains as well as the resources when integrated across the entire spatial habitat (Figures S5 and S8). Additionally we note the frequency dependence of motility selection in our model (Figure S7). We find that in addition to replicating the experimental results for our 4 treatments, the model also predicts motility selection in the monoculture case i.e. when the focal species is grown in isolation with one (or both) resources it requires for growth being distributed across the entire spatial habitat (Fig S4).

**Figure S4:**
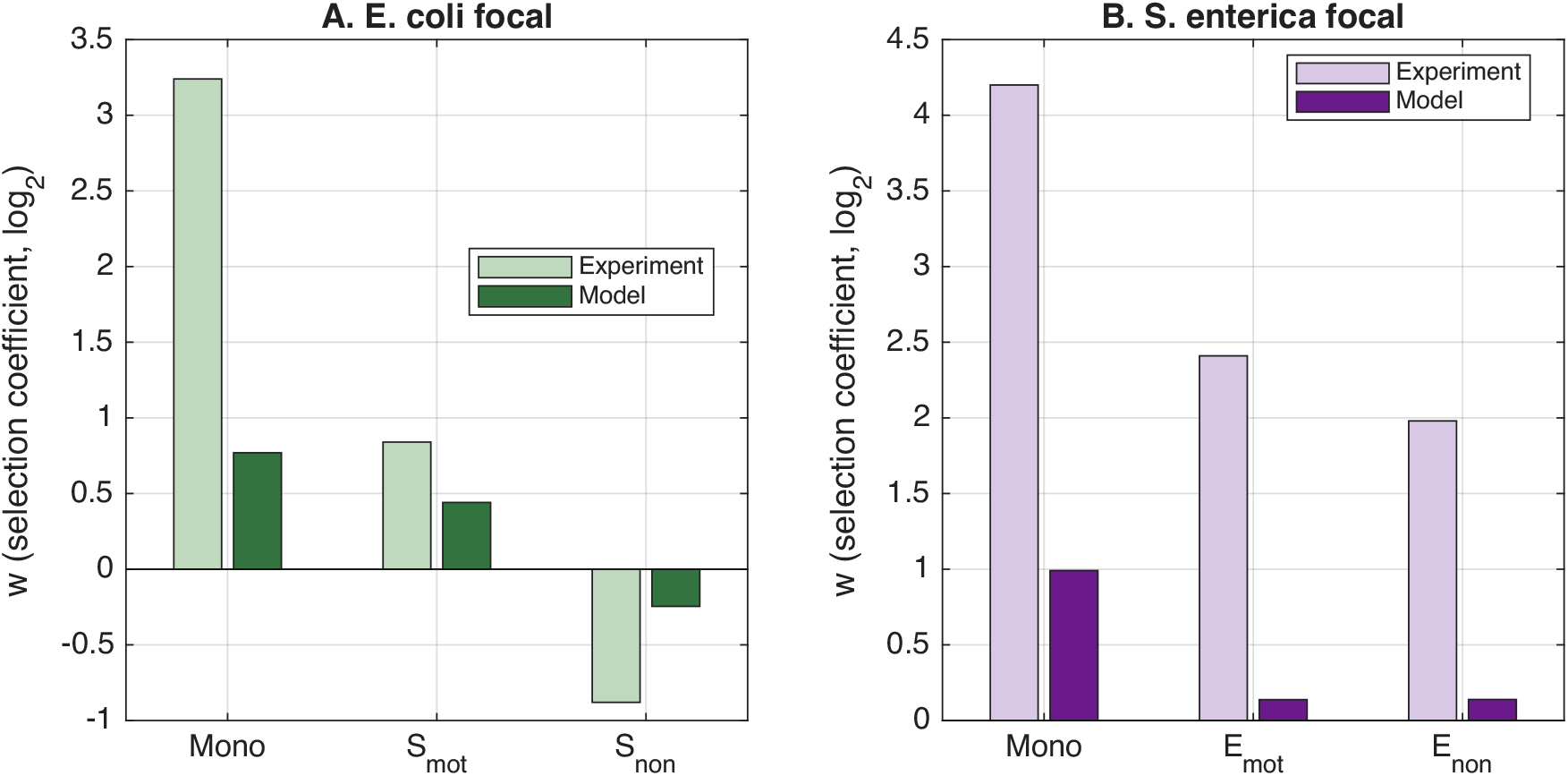
Motility is always selected for in monoculture. In panel (A), we find that in monoculture, the model predicts selection for motility in *E*.*coli*. We also replot results shown in **??**i where motility selection in treatments depend on partner motility status. In panel (B) similarly, we find that in monoculture as well as other treatments in co-culture, motility is always selected for.

#### S5.4 Supplementary figures associated with Figure 5

##### Generic qualitative match of simulations and experimental outcomes across large parameter ranges

In order to confirm that the results in Figure **??** showing a qualitative match with selection outcomes of experiments were not due to a carefully picked set of parameter values, we varied the accumulation ratios *ρ*_*M*_ and *ρ*_*A*_ across a wide range of values across an order of magnitude. We then calculated the fitness benefit of the motile strain at the end of the simulations i.e. ‘w’ (Figure S9a-d). We found that across a wide region of these values, where the mutualism did not break down (*ρ*_*M*_ *ρ*_*A*_ > 1), that the selection outcomes were similar to what we found in our experiments (S9e). This confirmed that our simulation results were not a result of handpicked parameters but an outcome of the interplay of underlying mechanisms in the model itself.

**Figure S5:**
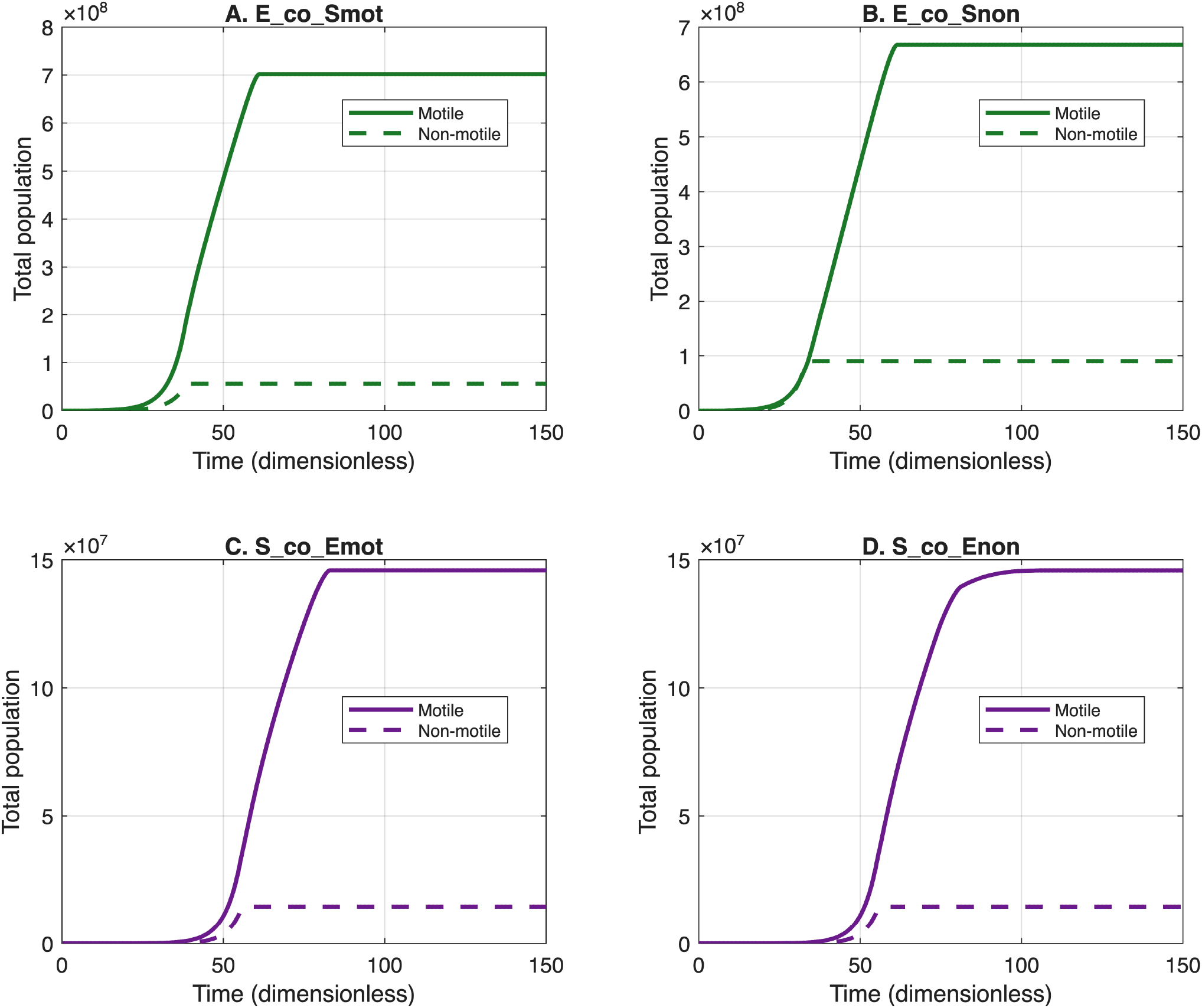
Population dynamics of focal strains in all treatments (summed across space) for all four co-culture treatments. All four treatments reach steady state and generically, *E. coli* grows to higher densities than *S. enterica* in all simulations due to higher maximal growth rates and greater amounts of resource accumulation i.e. *ρ*_*M*_ ≫ 1 and *ρ*_*A*_ ≪ 1. Note that the abundances here are shown after scaling. To obtain true abundances in the dimensional, unscaled model, mutliply densities by 10^6^. Although it appears that the motile genotype consistently appears to have higher density, note that its starting frequency is 0.9 compared to the non-motile strain which is 0.1.

**Figure S6:**
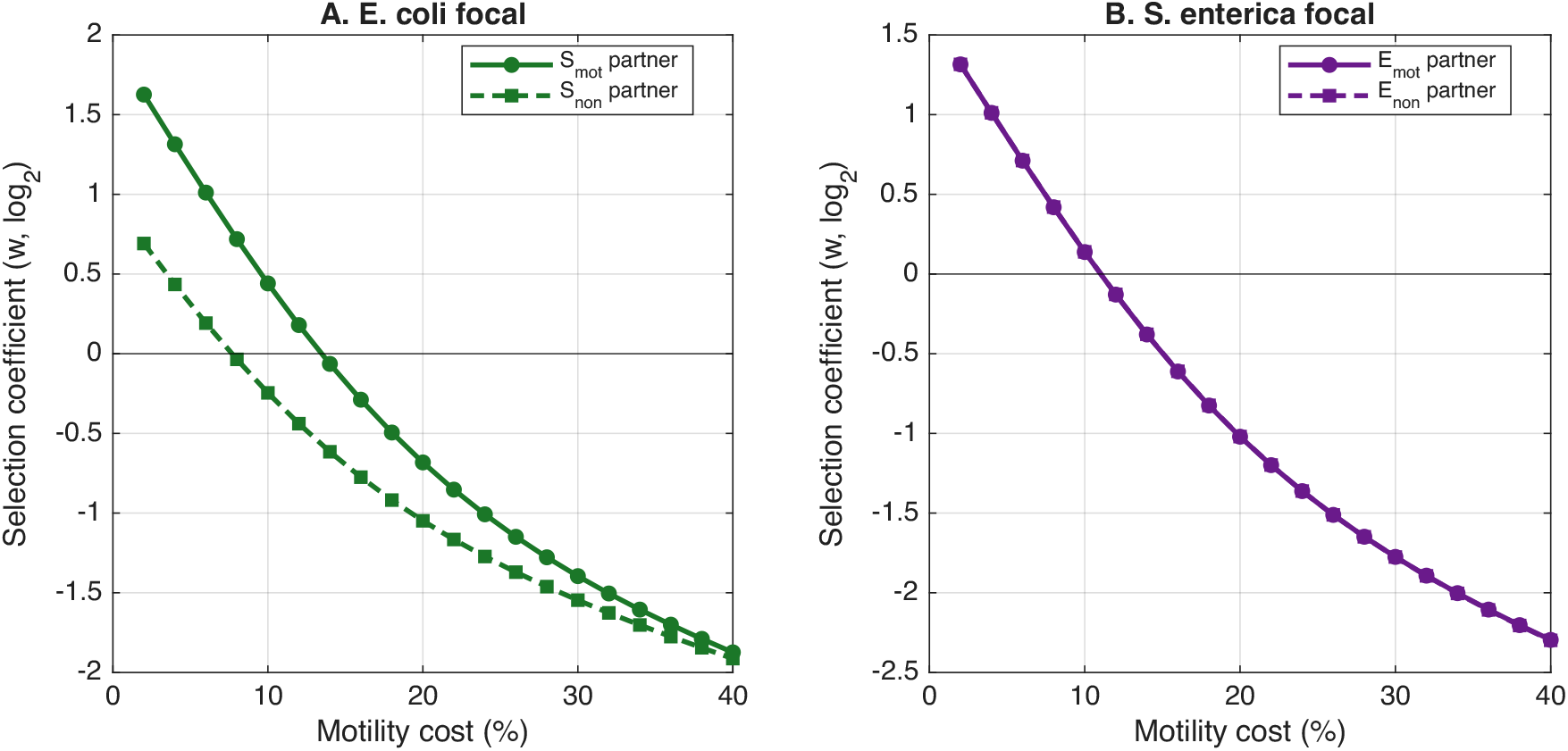
Increasing costs of motility alter selection outcomes and increase selection against motility. As motility costs are increased, panel (a) shows how selection against motility increases for *E. coli* strains for both motile and non-motile *S. enterica* partners. The transition to selection against motility occurs faster in the presence of a non-motile partner than a motile one. Panel (b) shows the same transition for *S. enterica* in the presence of different *E*.*coli* partners. Here, the cost as which transitions occur are much more similar between different partner motility statuses.

##### S5.4.1 Low motility cost and higher growth rate ratio select for motility across both species

Generic phase diagram emerges across different “treatments” wherein motility is always selected for at low motility costs which transitions to motility getting selected against with increasing costs. Motility is also selected for in S across a greater range of motility costs when its growth rate is higher relative to E i.e. *α* ≈ 1 or greater. Parameters used are the same as Figure 5 with *α* and c being varied (see Figure S10). Note that in all the figures in this section, the ‘star’ icon depicts the values “derived” from experimental values. All fixed parameters in this section are the same as the ones used in Figure 5.

In the main text, we focus on parameter regimes where lactose is the limiting resource. This is because limitation by either of the cross-fed resources implies the breakdown of the mutualistic association between the species whereas lactose limitation arises due to external resources being consumed entirely.

**Figure S7:**
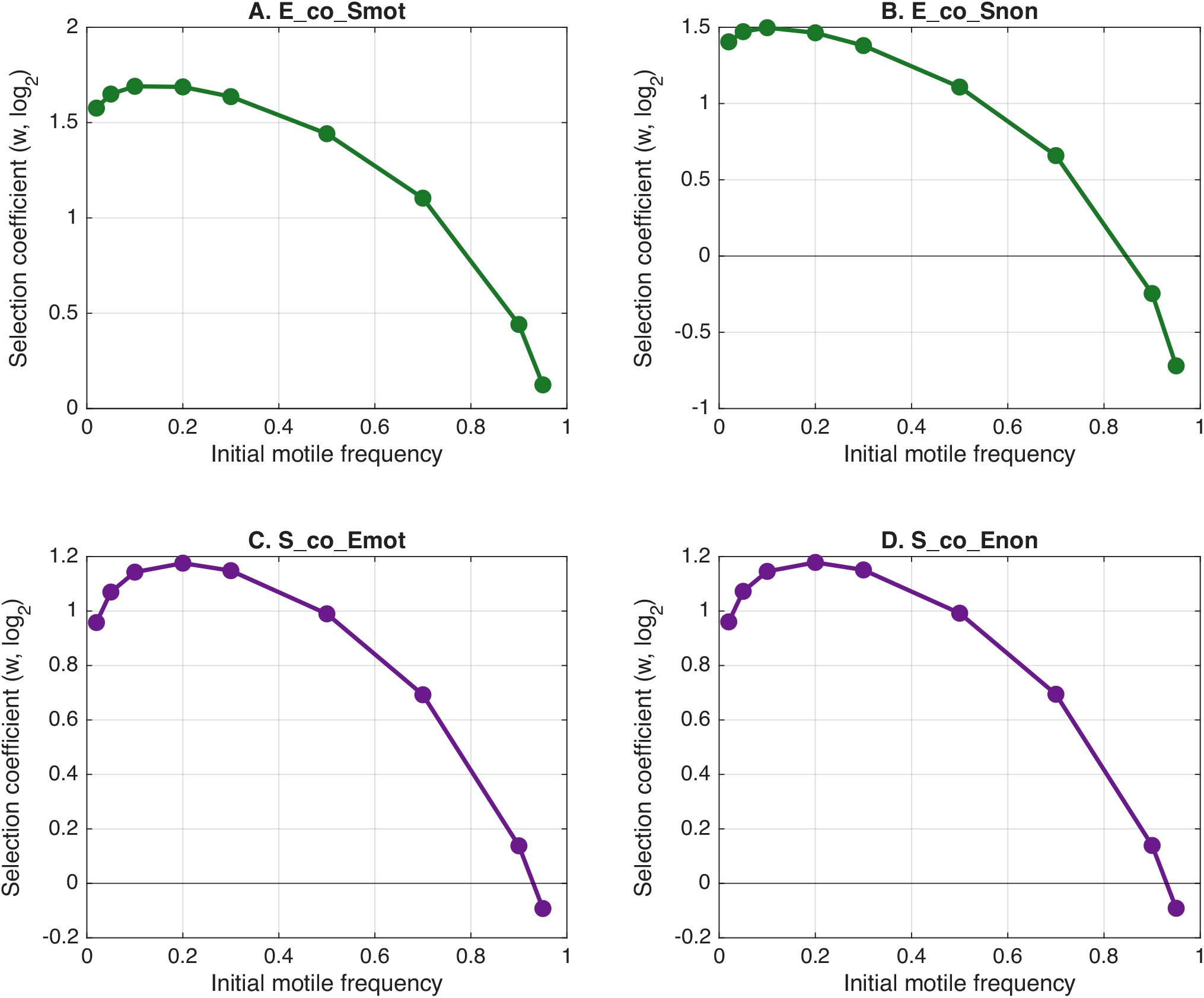
Frequency dependent strength of selection for motility is non-monotonic and generic across all treatments. All parameters other than initial density is fixed and the same as Figure **??** in the main text except initial ratio of motile and non-motile strains.

**Figure S8:**
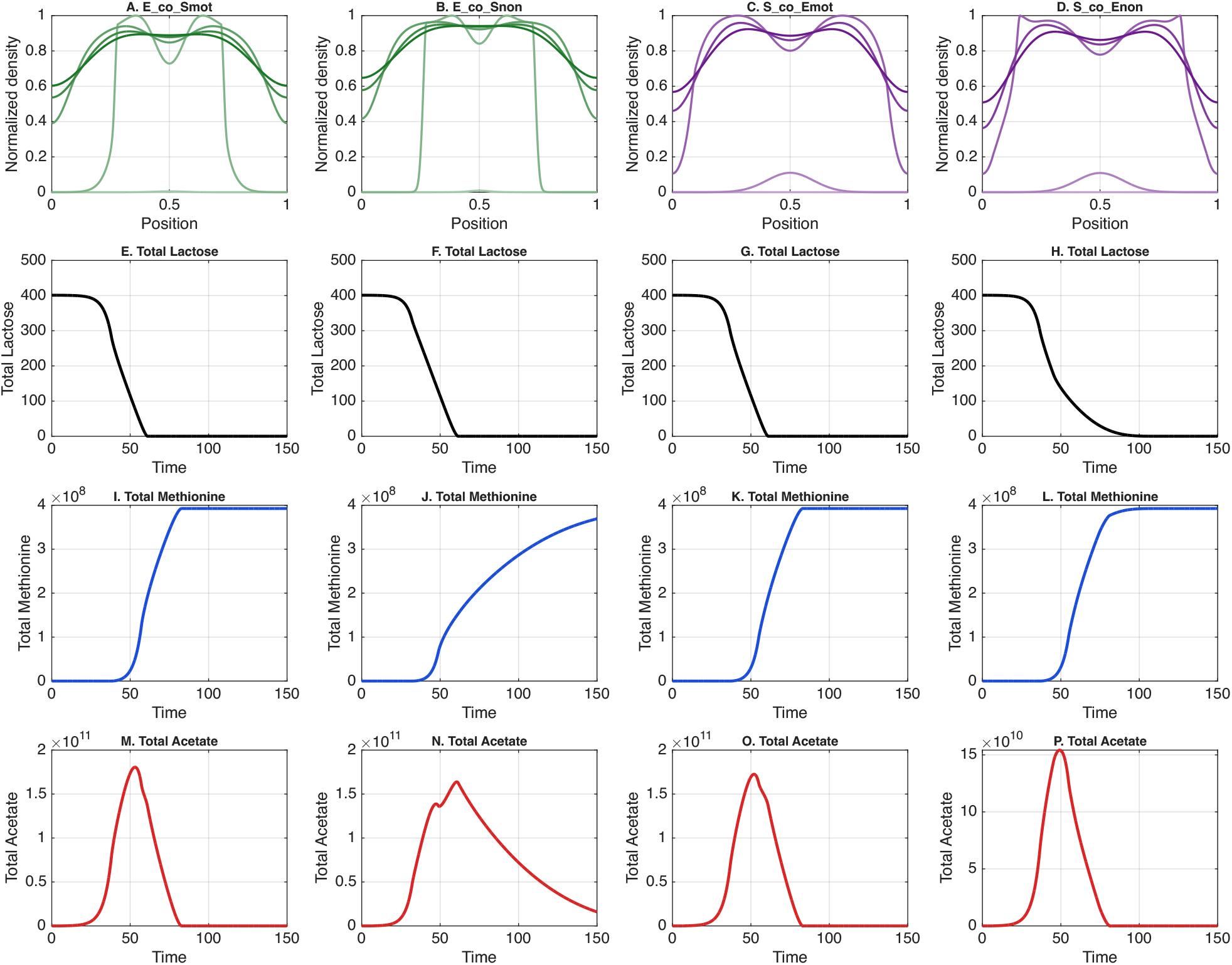
Temporal dynamics of bacteria and resource concentrations. We capture the temporal dynamics of the simulations performed in Figure **??** to establish equilibrium attainment and to visualise endpoint dynamics. In row 1 (panels A-D), we show the temporal abundance distribution of the motile strain of the focal species for each of the four treatments evaluated in the model. Increasingly darker and opaque curves depict later time points. In row 2 (panels E-H), we show the depletion of lactose in the system signalling the end of growth of the culture (thus the end of our simulations). Row 3 (panels I-L) shows the accumulation of methionine in our system levelling off at longer periods of time. This indicates once again that all the lactose has been consumed by bacteria and methionine by-products have accumulated in the system. In contrast, acetate initially accumulates (row 4, panels M-P)before being entirely consumed in the system. These differing equilibrial resource densities are driven by the accumulation ratios of the respective resources i.e. *ρ*_*M*_ and *ρ*_*A*_.

**Figure S9:**
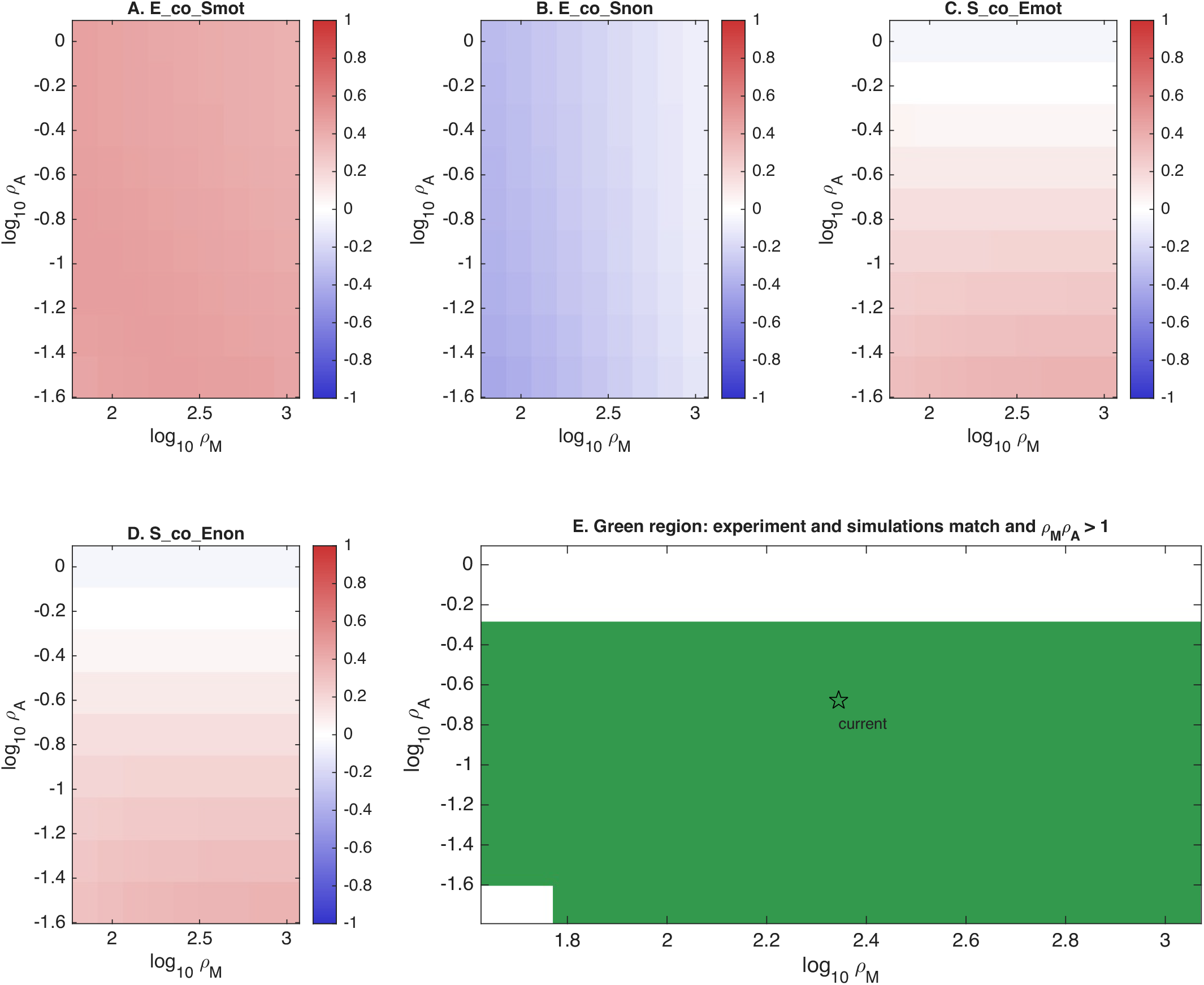
Large region of parameter space reproduces all four experimental outcomes. Panels A-D show selection on motility across methionine and acetate accumulation ratios for each of the four co-culture treatments. Colour indicates the selection coefficient *w* (log_2_). Panel E regions in green shows the combinations where all four selection coefficient signs match the experimental result and the mutual is self-sustaining (i.e. *ρ*_*M*_ *ρ*_*A*_ *>* 1; Eq. 11). The star marks the values derived from estimations from experiments (*ρ*_*M*_ = 221, *ρ*_*A*_ = 0.21), which lie within this region. All other parameters are at baseline (Table S2) and simulations at *N* = 401 with *c* = 0.10.

**Figure S10:**
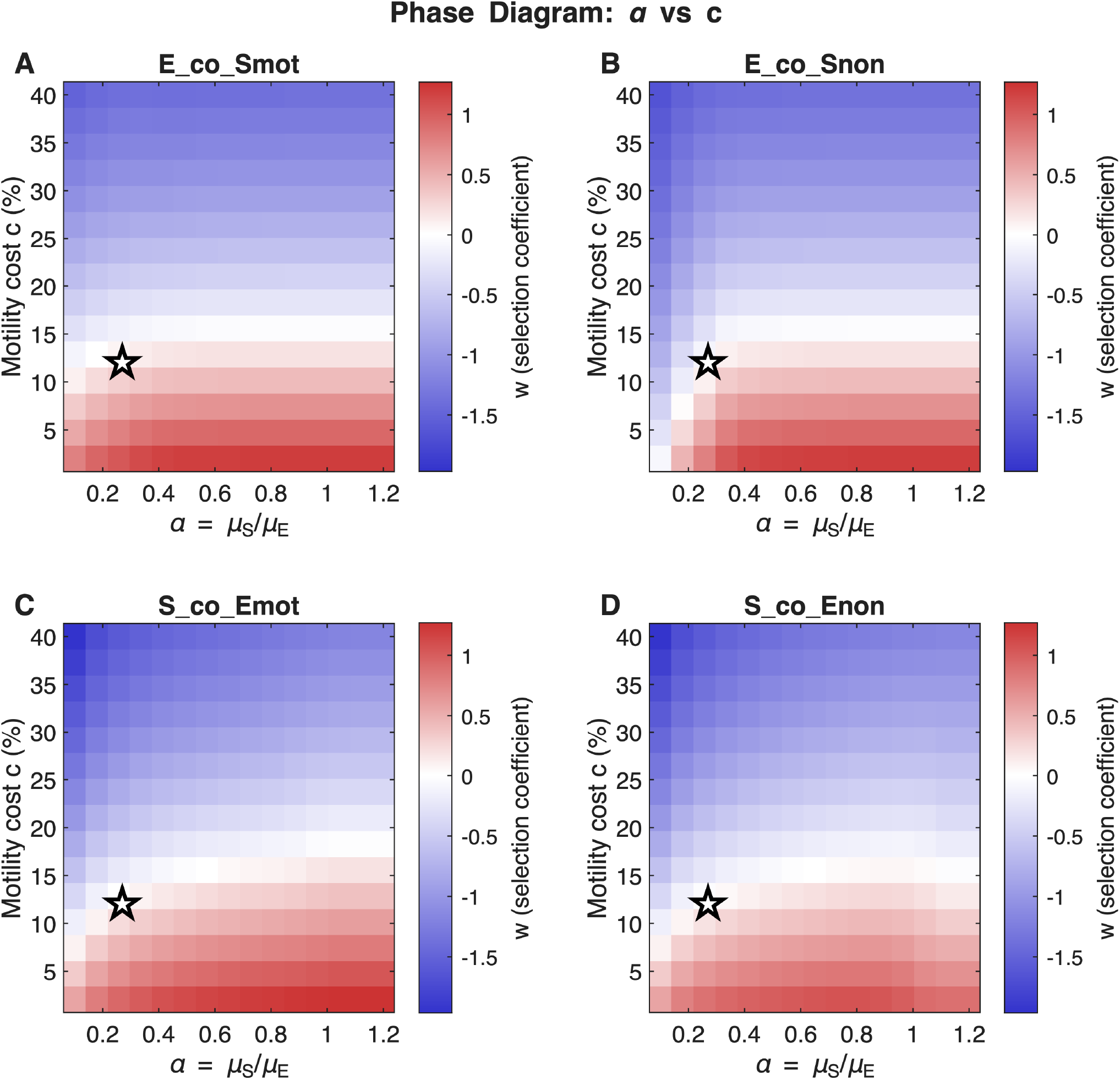
Motility selection phase diagram for varying costs of motility and ratio of maximal growth rates of species.

**Figure S11:**
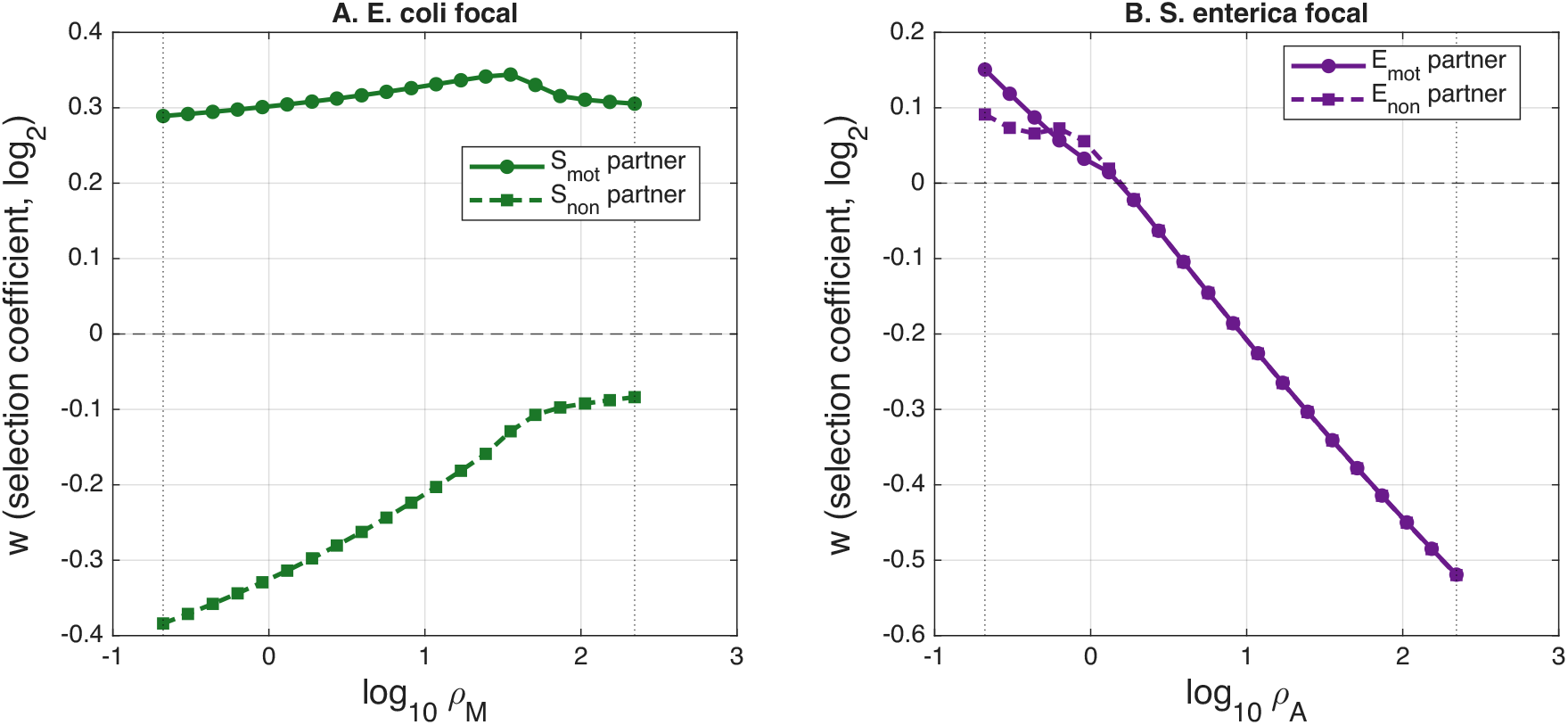
Differing selection outcome patterns with modified (flipped) *ρ* values. Selection transitions from selection for to against motility with increasing acetate production to consumption ratios for *S. enterica*. Selection signs do not change for *E. coli* due to resource co-limitation.

##### S5.4.2 Reversing metabolite accumulation ratios to verify joint role of partner growth and motility on selection

A numerical test of our mechanistic framework is to ask whether the asymmetry between species reverses when the metabolite dynamics are reversed. In our experimental system, methionine accumulates (*ρ*_*M*_ = 221 ≫ 1) while acetate is scarce (*ρ*_*A*_ = 0.21 < 1). Our framework predicts that if these ratios were reversed, for instance, by setting *ρ*_*M*_ = 0.21 (scarce methionine) and *ρ*_*A*_ = 221 (abundant acetate), the qualitative selection patterns should change in the following ways. First, *S. enterica* receives an abundant cross-fed metabolite (acetate as now *ρ*_*A*_ ≫ 1). Thus, acetate saturates the domain, eliminating the gradient that normally favours motility. According to our framework, because *S. enterica* consumes only one resource, it has no fallback gradient which can select for motility. Motility should therefore, in this case, be selected against regardless of partner status. Second, *E. coli* receives a scarce cross-fed metabolite in the flipped case (methionine, now *ρ*_*M*_ < 1). From our framework, we should expect methionine gradients should persist and motility should be favoured. However, there is a further complication because the methionine source is *S. enterica*, which grows slowly (*α* = 0.23). When *S. enterica* is non-motile, it barely expands from the inoculation point, creating an effectively static methionine source. Motile *E. coli* that disperses outward moves away from this static source into regions with lactose but no methionine. Because *E. coli* is co-limited by both methionine and lactose, it cannot grow without methionine, and motility becomes disadvantageous. When the partner is motile, however, methionine production is distributed and motile *E. coli* can access it.

Simulations with *ρ*_*M*_ = 0.21 and *ρ*_*A*_ = 221 (all other parameters at baseline) confirm these predictions (Figure S11A and B). We find that *S. enterica* motility is now selected against in both partner conditions (*w* < 0), consistent with the reversal predicted by our framework. However, *E. coli* selection remains partner-dependent i.e. motility is favoured with a motile partner (*w >* 0) but disfavoured with a non-motile partner (*w* < 0). Scanning the accumulation ratios continuously from the experimental values to the flipped values reveals smooth transitions in selection (Figure S11B). For *S. enterica*, the transition is clean i.e. *w* crosses zero near *ρ*_*A*_ ≈ 1 whereas for *E. coli*, the motile-partner curve remains positive (motility is selected for) across the entire range of *ρ*_*M*_ examined, while the non-motile-partner curve transitions from strongly negative (motility is selected against) to less negative (but never positive) with increasing *ρ*_*M*_ (Figure S11A).

The persistence of partner-dependent selection in *E. coli* across both the original and flipped regimes highlights the interaction between co-limitation and partner spatial structure. Because *E. coli* requires both lactose and methionine at the same location, the benefit of motility always depends on whether methionine is available at the colony periphery. This is partner dependent as a motile partner distributes methionine production while a non-motile, slow-growing partner keeps it localised. Whether methionine is globally abundant (original system, *ρ*_*M*_ = 221) or globally scarce (flipped system, *ρ*_*M*_ = 0.21), the fundamental constraint is that resource co-limited species are sensitive to the spatial distribution of their cross-fed metabolite in a way that single-resource species are not.

These results demonstrate that the species asymmetry in motility selection is not an intrinsic property of *E. coli* or *S. enterica*, but arises from the metabolic nature of the interaction. Any species receiving a single abundant metabolite (*S. enterica* in the flipped regime) will experience selection against motility. Any co-limited species whose partner is slow-growing and non-motile (*E. coli* in both regimes) will continue to experience partner-dependent selection. Thus, growth rate asymmetry between partners (*α*) plays a key role in determining how mobile the metabolite source is even in the absence of active motility. Next, we show that when we run a simulation experiment assuming *E. coli* growth only depends on methionine (i.e. single resource dependence), we find that selection for motility occurs regardless of partner motility status recapitulating the dynamics seen when *S. enterica* is the focal species (Fig S12).

**Figure S12:**
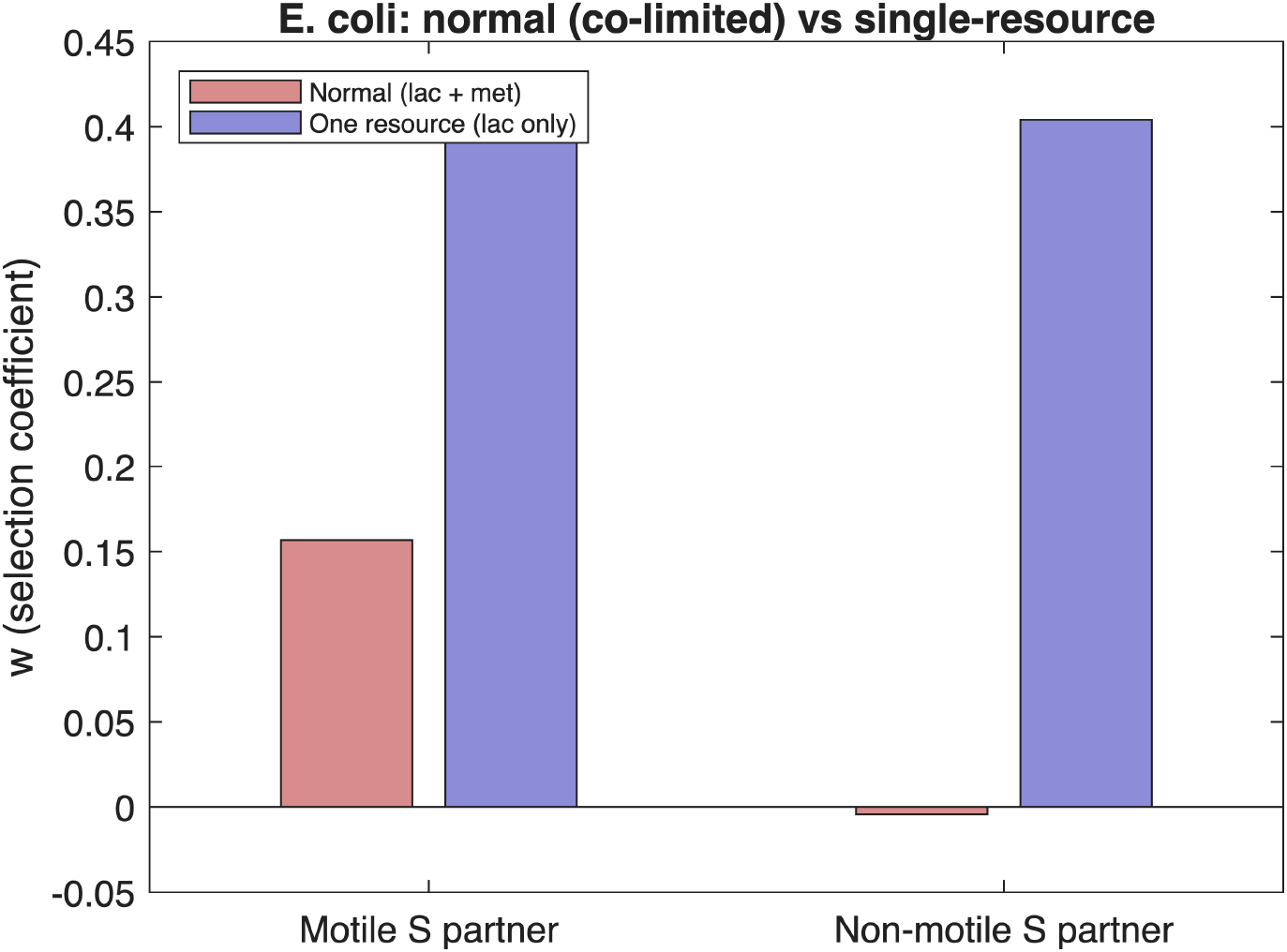
Simulations of a “synthetic” *E. coli* with single resource dependence (lactose) shows selection for motility regardless of partner motile status (blue) in contrast to true experimental scenario wherein *E. coli* depends on two resources for growth (red bar). Left bars show selection in the presence of motile partner and right bars show the presence of a non-motile partner.

## Notes

### Competing Interest Statement

The authors have declared no competing interest.

## References

[1] Hall-Stoodley L, Costerton JW, Stoodley P. Bacterial biofilms: from the Natural environment to infectious diseases. Nature Reviews Microbiology. 2004 Feb;2(2):95–108. Available from: https://www.nature.com/articles/nrmicro821.

[2] Davey ME, O’toole GA. Microbial Biofilms: from Ecology to Molecular Genetics. Microbiology and Molecular Biology Reviews. 2000 Dec;64(4):847–67. Available from: https://journals.asm.org/doi/abs/10.1128/mmbr.64.4.847-867.2000.

[3] Flemming HC, Wuertz S. Bacteria and archaea on Earth and their abundance in biofilms. Nature Reviews Microbiology. 2019 Apr;17(4):247–60. Available from: https://www.nature.com/articles/s41579-019-0158-9.

[4] Nadell CD, Drescher K, Foster KR. Spatial structure, cooperation and competition in biofilms. Nature Reviews Microbiology. 2016 Sep;14(9):589–600. Available from: https://www.nature.com/articles/nrmicro.2016.84.

[5] Yanni D, Márquez-Zacarías P, Yunker PJ, Ratcliff WC. Drivers of Spatial Structure in Social Microbial Communities. Current Biology. 2019 Jun;29(11):R545–50. Available from: https://www.cell.com/current-biology/abstract/S0960-9822(19)30396-3.

[6] de Vos MGJ, Zagorski M, McNally A, Bollenbach T. Interaction networks, ecological stability, and collective antibiotic tolerance in polymicrobial infections. Proceedings of the National Academy of Sciences. 2017 Oct;114(40):10666–71. Available from: https://www.pnas.org/doi/abs/10.1073/pnas.1713372114.

[7] Kong W, Meldgin DR, Collins JJ, Lu T. Designing microbial consortia with defined social interactions. Nature Chemical Biology. 2018;14(8):821–9.

[8] Estrela S, Libby E, Cleve JV, Débarre F, Deforet M, Harcombe WR, et al. Environmentally Mediated Social Dilemmas. Trends in Ecology & Evolution. 2019 Jan;34(1):6–18. Available from: https://www.cell.com/trends/ecology-evolution/abstract/S0169-5347(18)30249-0.

[9] Chacón JM, Hammarlund SP, Martinson JN, Smith Jr LB, Harcombe WR. The ecology and evolution of model microbial mutualisms. Annual Review of Ecology, Evolution, and Systematics. 2021;52(1):363–84.

[10] Ghoul M, Mitri S. The ecology and evolution of microbial competition. Trends in Microbiology. 2016;24(10):833–45.

[11] O’Brien S, Hodgson DJ, Buckling A. Social evolution of toxic metal bioremediation in Pseudomonas aeruginosa. Proceedings of the Royal Society B: Biological Sciences. 2014;281(1787).

[12] Jumpponen A, Trappe JM. Dark septate endophytes: a review of facultative biotrophic rootcolonizing fungi. New Phytologist. 1998 Oct;140(2):295–310. Available from: https://nph.onlinelibrary.wiley.com/doi/10.1046/j.1469-8137.1998.00265.x.

[13] Croft MT, Lawrence AD, Raux-Deery E, Warren MJ, Smith AG. Algae acquire vitamin B12 through a symbiotic relationship with bacteria. Nature. 2005 Nov;438(7064):90–3. Available from: https://www.nature.com/articles/nature04056.

[14] Dekas AE, Poretsky RS, Orphan VJ. Deep-Sea Archaea Fix and Share Nitrogen in Methane-Consuming Microbial Consortia. Science. 2009 Oct;326(5951):422–6. Available from: https://www.science.org/doi/abs/10.1126/science.1178223.

[15] Zelezniak A, Andrejev S, Ponomarova O, Mende DR, Bork P, Patil KR. Metabolic dependencies drive species co-occurrence in diverse microbial communities. Proceedings of the National Academy of Sciences. 2015 May;112(20):6449–54. Available from: https://www.pnas.org/doi/10.1073/pnas.1421834112.

[16] Wei Y, Wang X, Liu J, Nememan I, Singh AH, Weiss H, et al. The population dynamics of bacteria in physically structured habitats and the adaptive virtue of random motility. Proceedings of the National Academy of Sciences. 2011 Mar;108(10):4047–52. Available from: https://www.pnas.org/doi/abs/10.1073/pnas.1013499108.

[17] Cremer J, Honda T, Tang Y, Wong-Ng J, Vergassola M, Hwa T. Chemotaxis as a navigation strategy to boost range expansion. Nature. 2019 Nov;575(7784):658–63. Available from: https://www.nature.com/articles/s41586-019-1733-y.

[18] Kearns DB. A field guide to bacterial swarming motility. Nature Reviews Microbiology. 2010 Sep;8(9):634–44. Available from: https://www.nature.com/articles/nrmicro2405.

[19] Ni B, Colin R, Link H, Endres RG, Sourjik V. Growth-rate dependent resource investment in bacterial motile behavior quantitatively follows potential benefit of chemotaxis. Proceedings of the National Academy of Sciences. 2020 Jan;117(1):595–601. Available from: https://www.pnas.org/doi/abs/10.1073/pnas.1910849117.

[20] Colin R, Sourjik V. Emergent properties of bacterial chemotaxis pathway. Current Opinion in Microbiology. 2017;39:24–33.

[21] Colin R, Ni B, Laganenka L, Sourjik V. Multiple functions of flagellar motility and chemotaxis in bacterial physiology. FEMS Microbiology Reviews. 2021;45(6):fuab038.

[22] Fraebel DT, Mickalide H, Schnitkey D, Merritt J, Kuhlman TE, Kuehn S. Environment determines evolutionary trajectory in a constrained phenotypic space. eLife. 2017;6:e24669.

[23] Dal Co A, van Vliet S, Kiviet DJ, Schlegel S, Ackermann M. Short-range interactions govern the dynamics and functions of microbial communities. Nature Ecology & Evolution. 2020 Mar;4(3):366–75. Available from: https://www.nature.com/articles/s41559-019-1080-2.

[24] Philip JS, Grewal S, Scadden J, Puente-Lelievre C, Matzke NJ, McNally L, et al. Mapping the loss of flagellar motility across the tree of life. The ISME Journal. 2025;19(1):wraf111.

[25] Kubisch A, Holt RD, Poethke HJ, Fronhofer EA. Where am I and why? Synthesizing range biology and the eco-evolutionary dynamics of dispersal. Oikos. 2014;123(1):5–22.

[26] Mack KM. Selective feedback between dispersal distance and the stability of mutualism. Oikos. 2012;121(3):442–8.

[27] Chaianunporn T, Hovestadt T. Evolution of dispersal in metacommunities of interacting species. Journal of Evolutionary Biology. 2012;25(12):2511–25.

[28] Wang M, Schubert OT, Ackermann M. Cell motility enhances metabolic coupling in spatially structured microbial communities. bioRxiv. 2026:2026–04.

[29] Scarinci G, Sourjik V. Impact of direct physical association and motility on fitness of a synthetic interkingdom microbial community. The ISME Journal. 2023;17(3):371–81.

[30] Momeni B, Waite AJ, Shou W. Spatial self-organization favors heterotypic cooperation over cheating. elife. 2013;2:e00960.

[31] Harcombe WR, Riehl WJ, Dukovski I, Granger BR, Betts A, Lang AH, et al. Metabolic resource allocation in individual microbes determines ecosystem interactions and spatial dynamics. Cell reports. 2014;7(4):1104–15.

[32] Harcombe W. NOVEL COOPERATION EXPERIMENTALLY EVOLVED BETWEEN SPECIES. Evolution. 2010;64(7):2166–72. Available from: https://onlinelibrary.wiley.com/doi/abs/10.1111/j.1558-5646.2010.00959.x.

[33] Harcombe WR, Chacón JM, Adamowicz EM, Chubiz LM, Marx CJ. Evolution of bidirectional costly mutualism from byproduct consumption. Proceedings of the National Academy of Sciences. 2018;115(47):12000–4. Available from: https://www.pnas.org/doi/abs/10.1073/pnas.1810949115.

[34] Bernstein HC, Carlson RP. Design, construction, and characterization methodologies for synthetic microbial consortia. In: Engineering and Analyzing Multicellular Systems: Methods and Protocols. Springer; 2014. p. 49–68.

[35] Thomason LC, Costantino N, Court DL. E. coli genome manipulation by P1 transduction. Current protocols in molecular biology. 2007;79(1):1–17.

[36] Porwollik S, Santiviago CA, Cheng P, Long F, Desai P, Fredlund J, et al. Defined single-gene and multi-gene deletion mutant collections in Salmonella enterica sv Typhimurium. PloS one. 2014;9(7):e99820.

[37] Foster KR, Bell T. Competition, not cooperation, dominates interactions among culturable microbial species. Current biology. 2012;22(19):1845–50.

[38] Kehe J, Ortiz A, Kulesa A, Gore J, Blainey PC, Friedman J. Positive interactions are common among culturable bacteria. Science Advances. 2021;7(45):eabi7159.

[39] Rosazza T, Al-Tameemi Z, Rodriguez-Verdugo A. Microbial cross-feeding interactions reshape evolutionary trajectories of consumers by preserving motility. bioRxiv. 2026:2026–05.

[40] Rodriguez-R LM, Conrad RE, Viver T, Feistel DJ, Lindner BG, Venter SN, et al. An ANI gap within bacterial species that advances the definitions of intra-species units. mBio. 2024;15(1):e02696–23.

[41] Aggarwal D, Bellis KL, Blane B, De Goffau MC, Wagner J, Ng DY, et al. Large-scale characterisation of the nasal microbiome redefines Staphylococcus aureus colonisation status. Nature Communications. 2025;16(1):10415.

[42] Rosen MJ, Davison M, Bhaya D, Fisher DS. Fine-scale diversity and extensive recombination in a quasisexual bacterial population occupying a broad niche. Science. 2015;348(6238):1019–23.

[43] Goyal A, Chure G. Paradox of the Sub-Plankton: Plausible Mechanisms and Open Problems Underlying Strain-Level Diversity in Microbial Communities. Environmental Microbiology. 2025;27(4):e70094.

[44] Posfai A, Taillefumier T, Wingreen NS. Metabolic trade-offs promote diversity in a model ecosystem. Physical Review Letters. 2017;118(2):028103.

[45] Mori M, Zhang Z, Banaei-Esfahani A, Lalanne JB, Okano H, Collins BC, et al. From coarse to fine: the absolute Escherichia coli proteome under diverse growth conditions. Molecular Systems Biology. 2021;17(5):MSB20209536.

[46] Pearce MT, Agarwala A, Fisher DS. Stabilization of extensive fine-scale diversity by ecologically driven spatiotemporal chaos. Proceedings of the National Academy of Sciences. 2020;117(25):14572–83.

[47] Garud NR, Good BH, Hallatschek O, Pollard KS. Evolutionary dynamics of bacteria in the gut microbiome within and across hosts. PLoS Biology. 2019;17(1):e3000102.

[48] Al-Tameemi Z, Rodríguez-Verdugo A. Microbial diversification is maintained in an experimentally evolved synthetic community. mSystems. 2024;9(11):e01053–24.

[49] Takahashi Y, Yoshimura J, Morita S, Watanabe M. Negative frequency-dependent selection in female color polymorphism of a damselfly. Evolution. 2010;64(12):3620–8.

[50] Christie MR, McNickle GG, French RA, Blouin MS. Life history variation is maintained by fitness trade-offs and negative frequency-dependent selection. Proceedings of the National Academy of Sciences. 2018;115(17):4441–6.

[51] Gigord LD, Macnair MR, Smithson A. Negative frequency-dependent selection maintains a dramatic flower color polymorphism in the rewardless orchid Dactylorhiza sambucina (L.) Soo. Proceedings of the National Academy of Sciences. 2001;98(11):6253–5.

[52] Gude S, Pinçe E, Taute KM, Seinen AB, Shimizu TS, Tans SJ. Bacterial coexistence driven by motility and spatial competition. Nature. 2020;578(7796):588–92.

[53] Fukami T. Historical contingency in community assembly: integrating niches, species pools, and priority effects. Annual Review of Ecology, Evolution, and Systematics. 2015;46(1):1–23.

[54] Peay KG, Belisle M, Fukami T. Phylogenetic relatedness predicts priority effects in nectar yeast communities. Proceedings of the Royal Society B: Biological Sciences. 2012;279(1729):749–58.

[55] Moraïs S, Mizrahi I. Micro-scale spatial metagenomics opens a new era in microbiome ecology. Trends in Microbiology. 2026.

[56] Saarenpää S, Shalev O, Ashkenazy H, Carlos V, Lundberg DS, Weigel D, et al. Spatial metatranscriptomics resolves host–bacteria–fungi interactomes. Nature Biotechnology. 2024;42(9):1384–93.

[57] Narla AV, Cremer J, Hwa T. A traveling-wave solution for bacterial chemotaxis with growth. Proceedings of the National Academy of Sciences. 2021;118(48):e2105138118.

[58] Narayanan N, Lutz P, Shaw AK. Coexistence of coinvading species with mutualism and competition. Ecology. 2025;106(2):e70039.

[59] Narayanan N, Shaw AK. Mutualisms impact species’ range expansion speeds and spatial distributions. Ecology. 2024;105(1):e4171.

[60] Soto-Martin EC, Warnke I, Farquharson FM, Christodoulou M, Horgan G, Derrien M, et al. Vitamin biosynthesis by human gut butyrate-producing bacteria and cross-feeding in synthetic microbial communities. mBio. 2020;11(4):10–1128.

[61] Sathe RR, Paerl RW, Hazra AB. Exchange of vitamin B1 and its biosynthesis intermediates shapes the composition of synthetic microbial cocultures and reveals complexities of nutrient sharing. Journal of Bacteriology. 2022;204(4):e00503–21.

[62] Fazzino L, Anisman J, Chacón JM, Heineman RH, Harcombe WR. Lytic bacteriophage have diverse indirect effects in a synthetic cross-feeding community. The ISME Journal. 2020;14(1):123–34.

